# Synthetic Biology - Mapping the Patent Landscape

**DOI:** 10.1101/483826

**Authors:** Paul Oldham, Stephen Hall

## Abstract

This article presents the global patent landscape for synthetic biology as a new and emerging area of science and technology. The aim of the article is to provide an overview of the emergence of synthetic biology in the patent system and to contribute to future research by providing a high quality tagged core dataset with 7,424 first filings and 71,887 family members. This dataset is intended to assist with evidence based exploration of synthetic biology in the patent system and with advancing methods for the analysis of new and emerging areas of science and technology.

The starting point for the research is recognition that traditional methods of patent landscape analysis based on key word searches face limitations when addressing new and emerging areas of science and technology. Synthetic biology can be broadly described as involving the design, synthesis and assembly of biological parts, circuits, pathways, cells and genomes. As such synthetic biology can be understood as emerging from a combination of overlaps and convergences between existing fields and disciplines, such as biotechnology, genetic engineering, protein engineering and systems biology. More precisely, synthetic biology can be understood as the *synthetic phase* of molecular biology and genetic engineering. This presents the problem that key word strategies may radically overestimate activity because they involve terms that are widely used in underlying fields that are contributing to synthetic biology.

In response to this problem we combined anthropology, scientometrics and data science to map authors from scientific publications on synthetic biology into the international patent system. We mapped 10,816 authors into the international patent system and identified 2,450 authors of articles on synthetic biology who are also inventors in the period to December 2017. By combining this data with citation information and a baseline keyword strategy we are able to describe the global patent landscape for synthetic biology and the wider patent universe in which synthetic biology is situated.

This article describes the main features of the global landscape and provides the tagged dataset as a contribution to evidence based debate on intellectual property and synthetic biology and methodological development. We anticipate that the data will prove useful in informing international policy debates on synthetic biology under the United Nations Convention on Biological Diversity.

## Introduction

Synthetic Biology is an emerging area of scientific research directed to the design, synthesis and assembly of biological parts, circuits, pathways, cells and genomes for the production of a wide range of products [1–3]. In the United States, the European Union and a number of other countries, governments are increasingly investing in synthetic biology research projects, programmes and networks.

Synthetic biology is also associated with public and policy debates on the implications of this emerging field for science, society and the environment. Debates cover a spectrum of issues ranging from biosafety, risks of bioerror and bioterror, the ethics of creating new life forms, the implications of synthetic biology for developing countries, and the implications of environmental release for the conservation and sustainable use of biodiversity [4–7].

Intellectual property has featured prominently in the literature on issues such as the economics of synthetic biology, ethical questions around the ownership of synthetic components and synthetic life and the implications of intellectual property for developing countries [8–19]. Intellectual property is also an important focus of concern within the synthetic biology community with some advocating open source approaches as exemplified in the work of the BioBricks Foundation [20–22].

Quantitative analysis of patent activity for synthetic biology has been limited relative to the emergence of synthetic biology as an international field of research. Existing quantitative studies have mainly focused on the scientific landscape for synthetic biology using key word based approaches [18,23–25] while patent analysis has focused on the extension of key word approaches into the patent system [18,26]. A small number of other studies have focused on fine grained exploration of patent activity for specific areas of interest such as chemical production pathways and Threonine producing strains of E. coli [27,28].

The ability of researchers to conduct patent analysis at scale is affected by access to the full text of patent documents, access to data on a global scale and identifying appropriate methods for navigating a global patent information system consisting of over 100 million documents. As discussed by Smith et. al. 2017 methodological transparency is now a significant issue in patent landscape analysis in the life sciences with reproducibility a major factor [29]. In this article our aim is to contribute to greater reproducibility in patent landscape research by both clearly defining the method and making the patent data freely available for wider use. In particular, we provide a manually validated and tagged dataset as an aid to future research and methodological development in areas such as machine learning [30].

Our aim in this article is not to address social, economic, ethical and environmental questions directly but instead to focus on improving the evidence base for democratic deliberation on these wider questions. Specifically, we focus on providing reliable and verifiable patent data as a platform for improving understanding of the nature of patent activity in this field, deliberation about its implications, and advancing methods to assess emerging areas of science and technology.

## Methods

In previous work we used a simple definition of synthetic biology consisting of “ ‘synthetic biology’ or ‘synthetic genomics’ or ‘synthetic genome’ or ‘synthetic genomes’ “ to map the international scientific landscape for synthetic biology as an emerging field [23]. Based on a simple keyword search in Web of Science we found that, after stemming, 356 terms captured 99% of the universe of scientific publications for synthetic biology in the period to 2010. Figure 1 is reproduced from the earlier work and displays top ranking terms.

**Figure 1:**
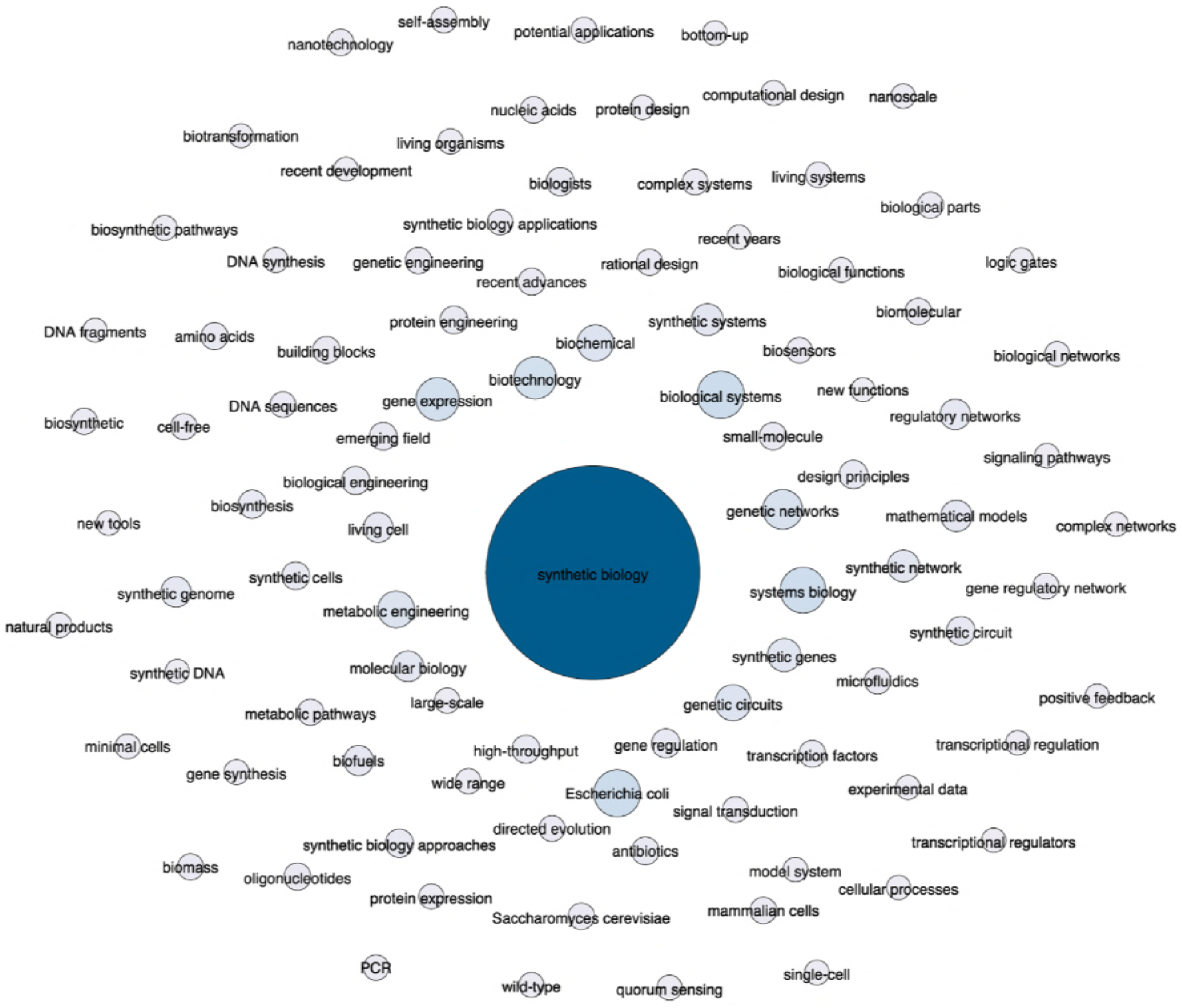
Top Terms in the Scientific Literature.

In considering the top terms we observed that “in the absence of a controlled vocabulary, the terms used in synthetic biology such as biotechnology or protein engineering are rapidly swamped by uses of the same terms in other research areas” [23].

There are two aspects to this problem. The first is whether it is reasonable to assume that fields related to synthetic biology that appear prominently in search results such as genetic engineering, protein engineering, systems biology, metabolic engineering or biological engineering should simply be subsumed within synthetic biology? The answer in our view must be no. The naive inclusion of such terms would lead to radical overestimation of an emerging field by including better established fields that are contributing to the emergence of synthetic biology. For example, Hu and Rousseau 2015 subsume “metabolic engineering”, “protein engineering” and “promoter engineering” within their definition of synthetic biology and in the process radically expand the landscape for synthetic biology (see below) [18]. Similar issues are observable in work by Van Doren et. al. 2013 who subsume systems biology, metabolic engineering and genome engineering within synthetic biology [26].

The problems associated with dependency on key words become particularly marked when we move into the 100 million patent documents in the global patent system. The first problem is data capture. Van Doren et. al. 2013 used the EPO World Patent Statistical Database to search the titles and abstracts of patent documents for synthetic biology related terms [26]. Patent documents consist of a title, abstract, description, claims and (in relevant cases) sequence listings and drawings. Searches of only the titles and abstracts will *de facto* underestimate activity in cases where a term of interest is only mentioned in the description or claims. This type of problem can readily be illustrated by carrying out a search for the term “genetic circuit” using the open access full text Lens patent database. A search of the title and abstracts in July 20107 revealed 66 documents in 36 families, a search of the titles, abstracts and claims revealed 82 results in 46 families while a search of the full text reveals 637 documents in 302 patent families. In this simple example, the search restricted to titles and abstracts captured 10% of the documents containing the term in the full text. Definitional exercises focusing only on titles and abstracts will underestimate activity because they ignore the major parts of patent texts.

However, a shift to searching patent full texts exposes the second aspect of the problem. It is common practice to develop keyword search strategies using the titles, abstracts and author keywords from the scientific literature and to deploy these terms in patent databases. Figure 2 presents the results of a search of the Web of Science Core Collection topic field in July 2017 for a set of terms commonly associated with synthetic biology. The topic field consists of titles, abstracts, author keywords and the titles of references (keywords plus). Figure 2 shows the results when the same terms were entered into a full text search using the open access Lens patent database.

**Figure 2:**
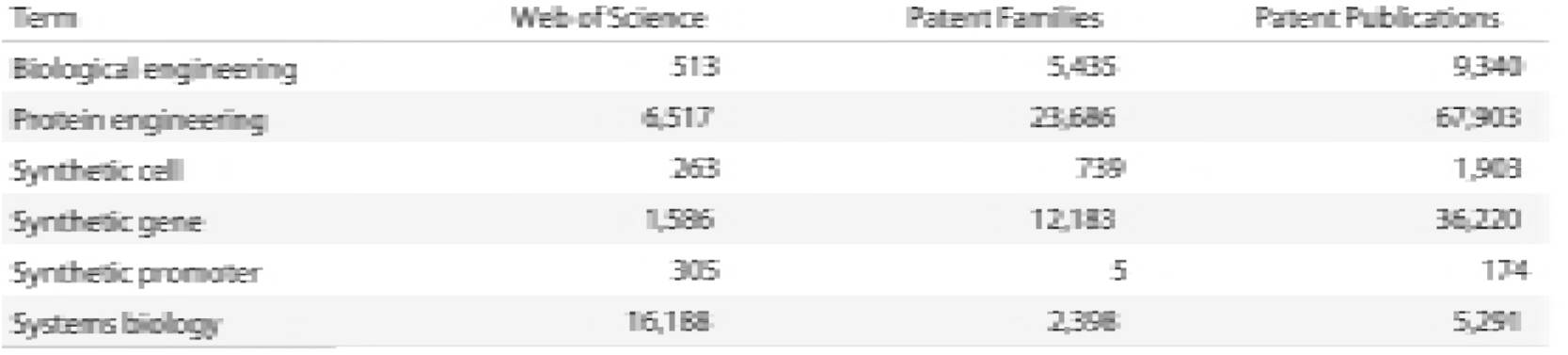
Web of Science Search vs. Patent Full Text Results 2017.

As we can readily observe, the consequences of moving terms such as protein engineering or synthetic gene into the patent system are dramatic. The divergence in the results immediately raises the question of whether the data refers to synthetic biology or alternative, and well established, alternate uses of terms in the patent system.

A common response to this type of problem is to use patent classification codes to restrict keyword searches to specific areas of the patent system. The patent system employs classification systems that were originally designed to assist examiners with retrieving documents and are now used for patent statistics. The main classification systems are the International Patent Classification (IPC) and the Cooperative Patent Classification (CPC). These are hierarchical classification systems consisting of alpha numeric codes that describe the content of patent documents at varying levels of detail from the Section (e.g. C = Chemistry; Metallurgy) down to the subgroup level such as C12Q1/68 for measuring/testing involving nucleic acids. From the 2000s onwards documents have normally been awarded multiple classification codes to more accurately describe their contents. Classification codes are frequently used for the elaboration of patent statistics and in bibliometrics/scientometrics. However, the use of classification codes to restrict searches of the patent system presents difficulties when working with emerging fields. First, while it is logical, as in the case of work by Van Doren et al (2013), to restrict patent searches to areas of the patent system concerned with biotechnology (e.g. class C12) in reality this assumes that which is to be demonstrated. As a field of research that places an emphasis on interdisciplinarity *a priori* restrictions run the risk of the analyst imposing a definition that does not reflect the actual characteristics of the emerging field.

As we will see in more detail below, in the specific case of synthetic biology a further problem is that viewed in terms of technical subject matter, rather than rhetorical description, synthetic biology is virtually indistinguishable from biotechnology and genetic engineering (e.g. IPC subclasses C12N, C12P, C12Q, CO7H etc.) [12]. In short, at present, the patent classification does not provide a ready means to distinguish synthetic biology from other areas of biotechnology within the patent system. As we discuss below, synthetic biology can perhaps best be understood as a phase shift *within* biotechnology and genetic engineering.

This is not an argument that keywords or combinations of keywords and classification schemes should be abandoned. As we will see below, keywords will continue to be a major part of the analysts tool kit. However, greater attention is required to the limitations of keywords approaches in circumstances where the full texts of 100 million patent documents in multiple languages are increasingly available for analysis. Continued reliance on title and abstract based definitions will provide insights but will underestimate activity. When applied in full text databases the same terms will frequently radically inflate activity.

In our earlier work we observed that “synthetic biology is a self-defining community of researchers from a variety of disciplines who are articulating themselves around the term synthetic biology and related terms such as synthetic genomics” [23]. Drawing on anthropology this suggested an alternative method focusing on “following the people” involved in synthetic biology rather than the language they speak [31]. Specifically, by mapping people (authors) from the scientific literature into the patent literature we would be in a position, to borrow a famous phrase from Bronislaw Malinowski, to let “the natives speak for themselves” rather than imposing analysts definitions [32].

The essence of this approach is to use a simple keyword search involving terms that are definitely distinctive to the emerging field as the seed for the identification of researchers working in the field. Based on a manually validated reference dataset we conducted the research in three phases.

In the first phase, we performed the simple baseline query for synthetic biology from our earlier work in mid-2014 in Web of Science (core collection) and retrieved data on 2,475 publications in the period to the end of 2013. The 2,475 publications contained a raw total of 6,224 raw author full names. After name cleaning and the exclusion of researchers in the social sciences, humanities and journalists, this produced a raw list of 5,379 author full names for use in searching the EPO World Patent Statistical Database (PATSTAT, October 2013 edition).

The PATSTAT person table (tls206_person) included 30,315,495 inventor names linked to 76,594,275 patent applications worldwide between 1782 and October 2013. From our starting point of 5,379 author full names we identified 3,160 raw name hits in PATSTAT. Following a period of experimentation with edit distances and Jaccard weightings, we established that an approach consisting of straightforward boolean queries that skipped punctuation worked best for achieving data capture. Validation of author name to inventor matching was facilitated by the reference set.

The mapping of inventor names is known as the ‘names game’ in patent statistics [33]. In the wider literature on the disambiguation of person names two important problems have been identified [34]. The first is the splitting of the same person name (synonyms) into multiple names through variations in punctuation or name reversals. This can be characterised as the “James T Kirk” problem where the same name could be expressed as “Kirk T James”, “James Kirk T” or “Kirk James T” with punctuation such as stops and commas introducing additional variations. It is therefore important to clarify whether the name variations denote the same person. The second, and more significant, problem is lumping (homonyms) where the same name may designate multiple persons [34]. Examples include John Smith in English, Carlos Garcia in Spanish, or Wei Wang in the case of China. In practice our research revealed that the lumping together of names of distinct persons is a far more difficult problem to address than name splitting. For example, in the case of the name Wei Wang we retrieved 3,855 patent records (applications) for this name. The scale of these results suggested either that Wei Wang is a super inventor or that the data involves multiple persons who have been lumped together on the same name.

The patent system also employs a citation system that is an important focus of economic research [35–37]. Specifically, citations form networks of documents that share one or more technical feature. We therefore surmised that data capture could be improved by reviewing cited and citing patent documents. However, we also recognised that citations may miss recent documents, such as those that have yet to be searched or examined, and potential outliers. For this reason keywords would play a continuing role in enhancing data capture. This formed the basis for a concentric circle model with 5 elements.

1. Precise. An author appears as an inventor on a patent document with a co-author of the same article.
2. Intersection. An author name co-occurs as an inventor on a document with the name of another author in the wider dataset.
3. Affiliation. An author name appears on a patent document with a match between the applicant name and the author affiliation field from Web of Science.
4. Citations. Citations of earlier patent documents (cited) and later patent filings (citing). Any names meeting criteria 1-3 were reallocated to the core set.
5. Keywords. The baseline query run on the full texts of patent collections worldwide with the aim of improving capture, particularly for recent developments.

The essence of the concentric model is to move from precision in the identification of records to completeness in the construction of a patent landscape. However, name co-occurrence based strategies confront particular challenges in East Asian countries, specifically China, where naming traditions lead to many shared names [38]. Our original tests revealed that naive co-occurrence based analysis with Chinese inventor names will result in the lumping together of networks of common names involving distinct and unrelated persons. The development of inventor name disambiguation algorithms is a growing field of research that has mainly focused on US data but with increasing attention to East Asian names [38–40]. During phase one we were unable to identify a method for addressing Chinese names in which we felt confident. However, we observed that of a total of 22,363 filings in China where author and inventor names intersected, 22,023 (98%) were filed only in China and 340 outside China. Based on this finding we restricted data for China to records where there was also a filing outside China. The practical consequence of this is that only international level documents originating from China were reviewed during this phase.

In the second phase we updated the Web of Science baseline data to mid 2017 and identified a total of 11,732 raw author names in 4,463 publications in Web of Science. Name cleaning was performed in VantagePoint from Search Technology Inc using author affiliations as the main match criteria. This led to 11,228 cleaned names. A total of 412 author names were identified for the social sciences, humanities and journalism and were excluded. A total of 10,816 researcher names were used to map across to the patent data.

Using the phase one dataset as a resource, all patent documents citing the original core set were retrieved and manually reviewed using the 10,816 names. The raw citing dataset until December 2017 consisted of 95,857 documents in 38,293 families. An updated keyword dataset was created by searching the full texts of patent databases world wide in Derwent Innovation Professional from Clarivate Analytics. The professional version of Derwent Innovation provides access to a wider range of full text collections (notably for Asia and the wider Americas) compared with other free or commercial patent databases. Author to inventor matching was then performed on the keyword set.

Each individual dataset was manually reviewed in VantagePoint by combining the Web of Science dataset with the corresponding patent dataset and creating a unified author full name and inventor field. We then used match criteria to group and label the dataset.

1. Coauthor. The author of an article appears with another author of the same article as a co-inventor on a patent document.
2. Intersection. The author of an article occurs as an inventor with the name of another author from the wider Web of Science data. This assists with capturing cases where researchers move over time.
3. Affiliation. A match between the name of an author as inventor and the affiliation in the publication. In the case of citation data (below) where relevant this included web lookup of the career history of researchers to identify former affiliations.
4. Subject Matter. In the absence of other match criteria we asked the following question: Does the subject matter of the patent application very closely match the subject matter of a scientific article by the author?
5. Other. In uncertain cases authors were looked up on the web. In cases where a coauthor in laboratory publication records appeared as an inventor with the focus author they were allocated to this group. In other cases authors who had moved organisation or founded spin off companies were allocated to this group.

The review process involved classifying documents as keep, review or exclude and recording the match criteria for each record. In cases where an element of uncertainty persisted around a retained record, the record was marked uncertain as an aid to methodological transparency.

The patent data was processed in VantagePoint and post processing and visualisation was performed in R, Gephi and Tableau [41]. Processing in VantagePoint consisted of removing duplicated publication records onto the first family member. Authors who were also inventors were combined into a single thesaurus during name cleaning. Inventors names outside the author group underwent basic name cleaning using shared priority filing numbers followed by shared applicant names as the clustering criteria. The cleaning process used fuzzy matching logic and each grouping was manually reviewed. Applicant cleaning was performed using the applicant name and Derwent applicant name codes and then further cleaned using fuzzy logic matching. The original applicant names as they appear on the public records and cleaned author names are provided in the dataset. All applicants in the core set were matched to a Company, Academic, Government, Hospital category while noting that in some cases applicant categories can be ambiguous. The International Patent Classification field was used to identify the main technology sectors. Text mining was performed in VantagePoint on the words and phrases from the titles, abstracts and claims. The focus of text mining was on identifying topics of interest to the community, such as promoters, reporters, CRISPR/Cas9 etc. and the uses of the claimed invention. Processing in R was performed using the popular tidyverse packages and the complete R code used for calculations is provided in the supplementary material [42].

Patent documents are by definition public data and the patent datasets are provided in simple.csv files containing the public patent data fields. Derwent Innovation includes a wide range of additional value added data fields beyond those that are part of the public file record. These fields are excluded from the dataset. Patent databases vary in their document number formats and we provide a standardised number that should work across multiple databases such as esp@cenet, PATSTAT, the Lens and WIPO Patentscope.

In approaching the data presented in this article it is important to note that patent analytics uses a range of patent counts and terminology. This can be confusing and in some cases misleading. In this article we use the following terminology.

1. The term patent document means either a patent application or a patent grant.
2. Patent family. Patent documents live in families that share one or more original filings as their parent or ‘priority’. When a patent application is filed for the first time anywhere in the world it becomes the ‘priority’ or first filing under the 1883 Paris Convention [43]. That priority filing may become the basis for additional applications and grants in multiple countries. In this article counts by patent families refer to counts of the very first filing, the document with the earliest priority date, in a set of related documents. Counts of first filings are therefore counts of patent families and we use these terms interchangeably. Because first filings are often not published we use the first available publication number as the representative of the first filing to enable the record to be looked up. The country code and date of this document should not be used for statistical purposes as they will be later than the first filing.
3. Family Members. Patent documents published anywhere in the world that share the same parent or priority are family members. The data on family members in this article uses the INPADOC definition of patent families (available in PATSTAT, Derwent Innovation, esp@cenet and the Lens). The INPADOC family definition also includes documents that, in the view of the examiners, are technically related and belong in the family even where they do not share an identical parent [44]. In a small number of cases for India, Singapore and Brazil no link has yet been established between a first filing and an INPADOC family. Minor adjustments were made in these cases to add the first publication as the family representative but in some cases these documents may be silently dropped from the analysis using family members.

## Results

The core landscape for synthetic biology consists of 7,424 families of patent applications and 71,887 family members worldwide in the period to December 2017. These patent families are situated in a wider patent universe consisting of 56,789 patent documents cited by the core landscape (cited or back citations) and 33,889 families with 499,284 family members that cite the core landscape (citing or forward citations) in the period to December 2017.

In total we identified 2,450 authors of articles on synthetic biology who are also inventors, representing approximately 23% of the 10,816 researchers publishing scientific research on synthetic biology. As this makes clear, the majority of researchers using the synthetic biology terms in our baseline literature dataset are not engaged in patent activity.

Figure 3 displays the overall trend in the first filings of new patent applications for the core dataset. The data has been reduced to the earliest document for each patent family to avoid duplication and is the date closest to the investment in research and development that led to the claimed invention. A linear model with Loess smoothing is used to establish the trend line. Figure 3 reveals a long tail of patent activity by researchers publishing in synthetic biology. Data is displayed until 2015 after which patent data displays a characteristic data cliff arising from lag times between the filing and publication of applications (typically between two and three years) (supplementary material).

**Figure 3:**
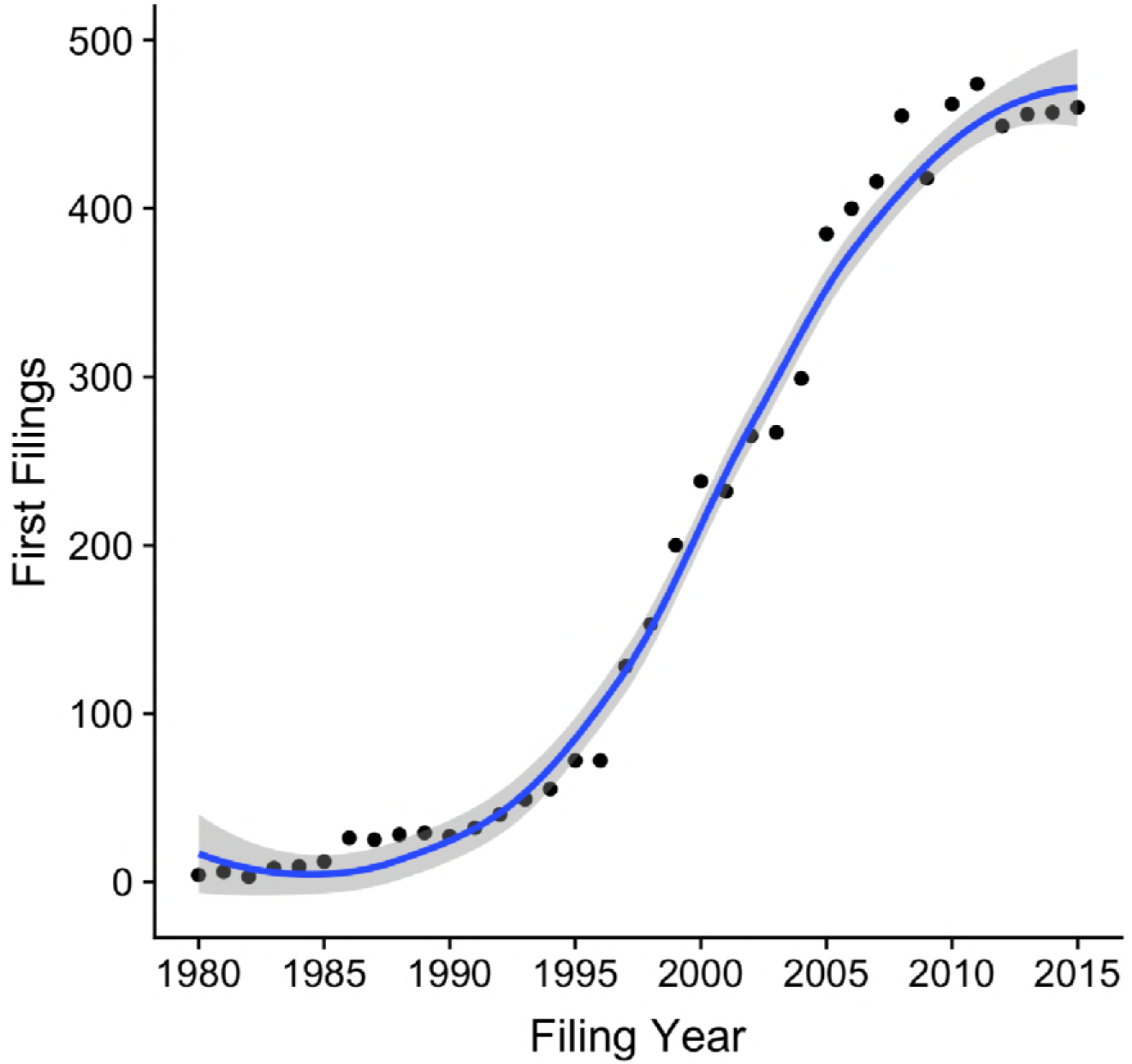
Trends in First Filings of Patent Families in the Core Landscape.

Figure 4 displays the components of the core landscape based on the main criteria used for the construction of the landscape. In Figure 4A documents in the set may fall into multiple groups. Figure 4B assigns the records appearing in more than one group to unique groups by privileging coauthors, affiliation, other (generally former affiliations), subject and keywords in that order.

**Figure 4:**
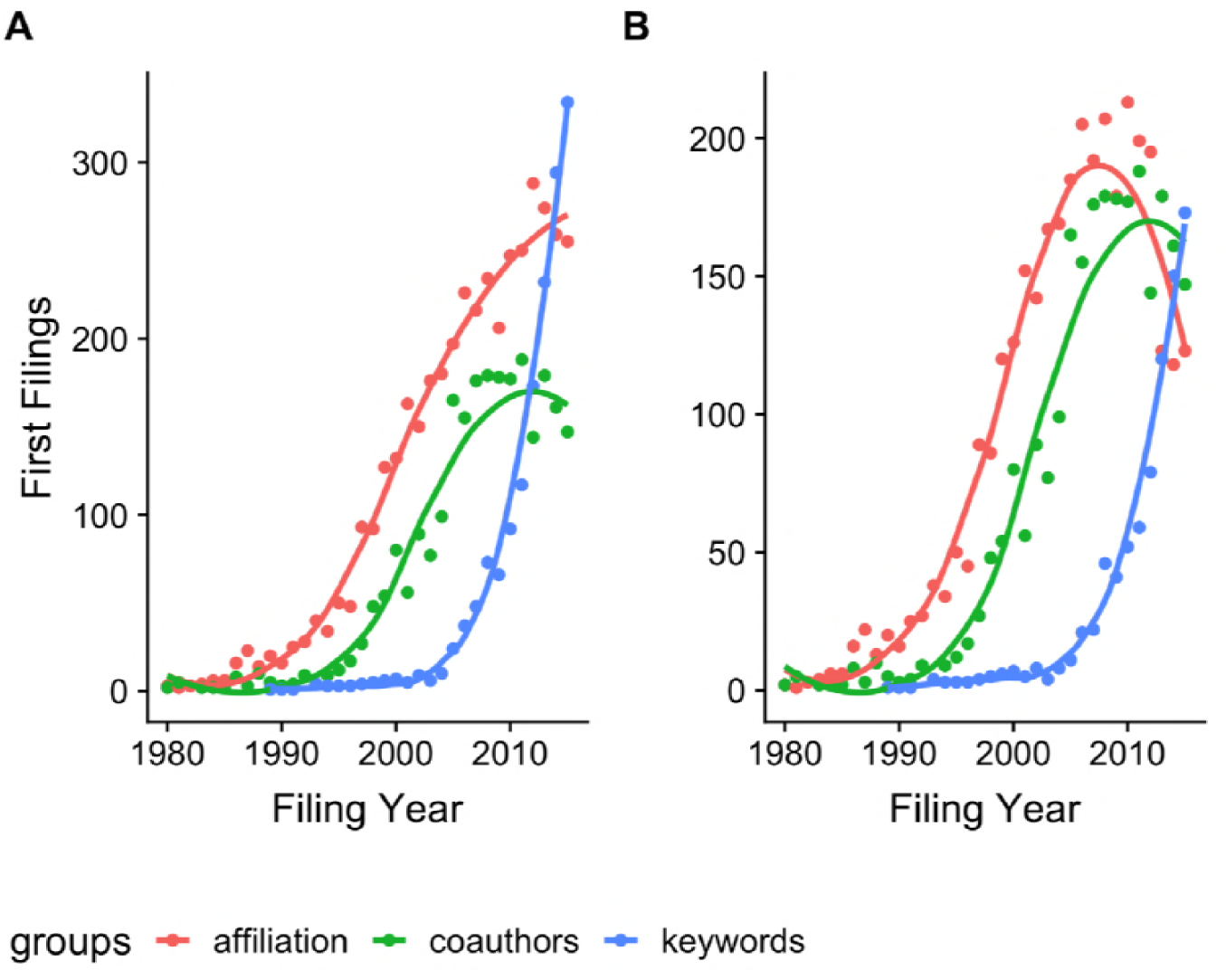
Core Landscape by Groups.

In considering Figure 4A we can clearly see that affiliation based matching proved to be the dominant criteria in the identification of patent documents linked to authors followed by co-authorship. We had anticipated that as synthetic biology becomes increasingly institutionalised in the form of laboratories, centres and programmes that coauthorship would logically emerge strongly as the dominant match criteria. That is, we expected that networks would emerge in patent activity involving synthetic biologists as the field consolidated and expanded. This appears to be confirmed by reweighting the groups to privilege coauthorship in Figure 4B with affiliation based matching being overtaken by coauthorship from approximately 2010 onwards. The extent to which network clustering effects will be reinforced in future is an open question. We examine inventor networks in detail below.

The most striking feature in Figure 4 is that the baseline keyword strategy performed poorly compared with coauthorship and affiliation based matching. For the majority of the period covered researchers publishing in synthetic biology did not use our simple baseline terms on synthetic biology, synthetic genomics or synthetic genome(s) anywhere in the full texts of their patent applications. While our keyword strategy was deliberately simple this exposes the wider difficulties involved in developing effective keyword strategies that do not radically under or over estimate activity. However, the ongoing importance of keyword based strategies is also revealed by the steepness of the curve as we move closer to the present. The steeply rising curve reflects the growing use of the baseline terms within the full texts of patent applications in the period between 2010 and 2015. In particular, closer examination revealed the increasing use of these terms in the texts of national patent filings in China. Thus, 457 (25%) of 1837 first filings in the dataset containing the keywords were from China. As noted above, our coauthor based approach focused only on applications from China that were also filed internationally. However, the growing availability of translations of Chinese patent documents meant that the baseline keyword strategy picked up an increasing number of Chinese inventors filing only in China.

Patent activity in synthetic biology as an emerging field is situated in a wider universe that can be mapped using patent citations. Citations of earlier patents (back citations or cited documents) that are relevant to the claimed invention are cited by applicants or examiners and form part of the public record. Patent activity by researchers in synthetic biology also has impacts on other patent applicants by limiting the scope of what later applicants may claim to be new or novel and involving an inventive step. This data is accessible through forward citations or citing documents. Taken together the cited, core and citing patent landscapes captures the majority of the patent universe in which synthetic biology is situated and evolving with keywords capturing additional recent activity and outliers.

Figure 5 displays trends in filings within the cited, core and citing landscapes based on the earliest filing date for each patent family.

**Figure 5:**
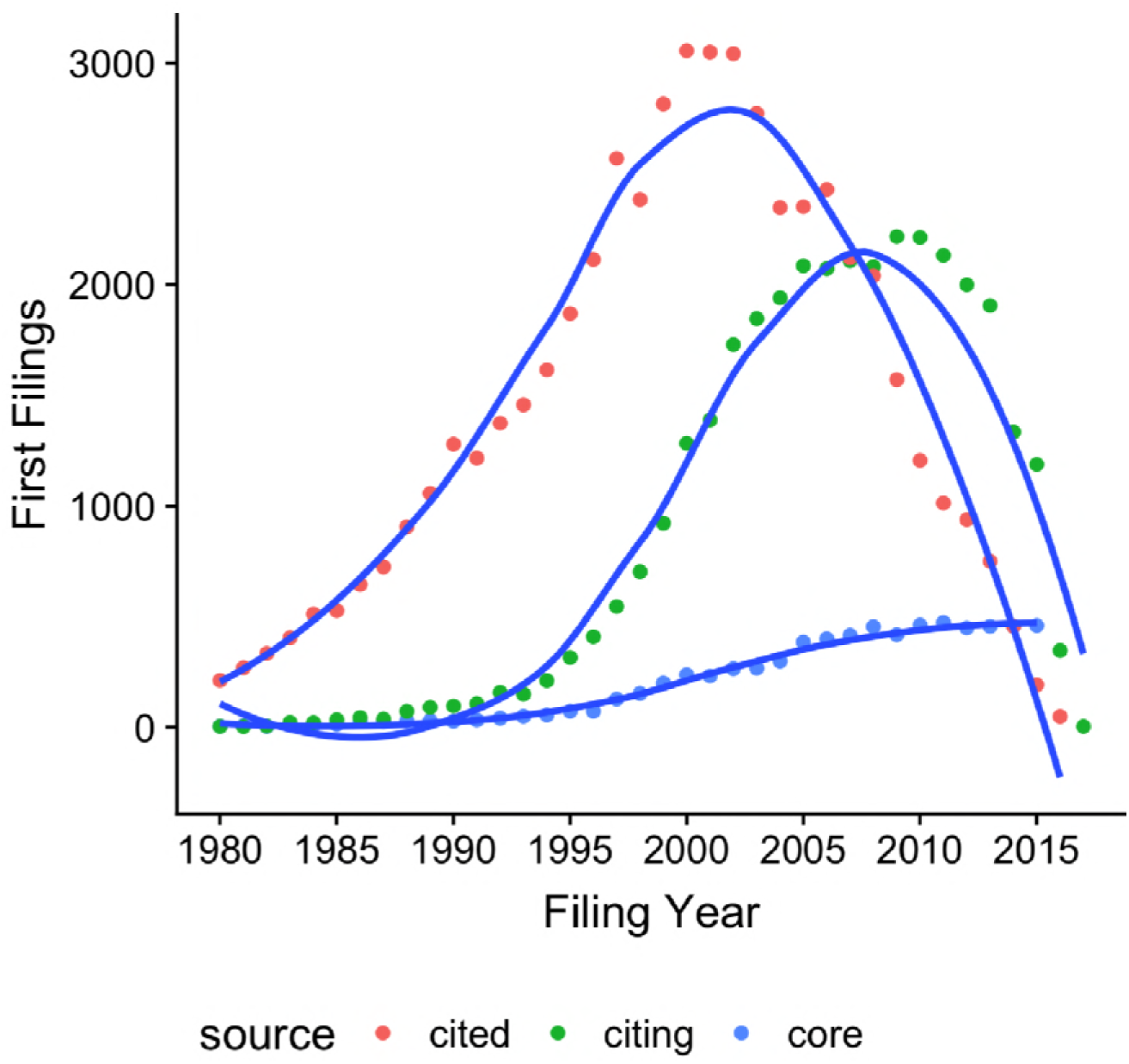
The Patent Universe for Synthetic Biology.

Figure 5 serves to demonstrate that synthetic biology within the patent system should not necessarily be seen as radical and “new” but will, to borrow from the American economist Suzanne Schotchmer, be cumulative in nature and stand on the shoulders of earlier giants [45]. These giants will be represented in the cited landscape and take us to the core of the emergence of biotechnology and genetic engineering in the patent system. Figure 6 displays the top ten cited patent documents in the core landscape ranked on the total citation scores in the patent system. The two top ranking documents are foundational for Polymerase Chain Reaction (PCR) technology while the directed evolution of proteins also features prominently in the cited landscape.

**Figure 6:**
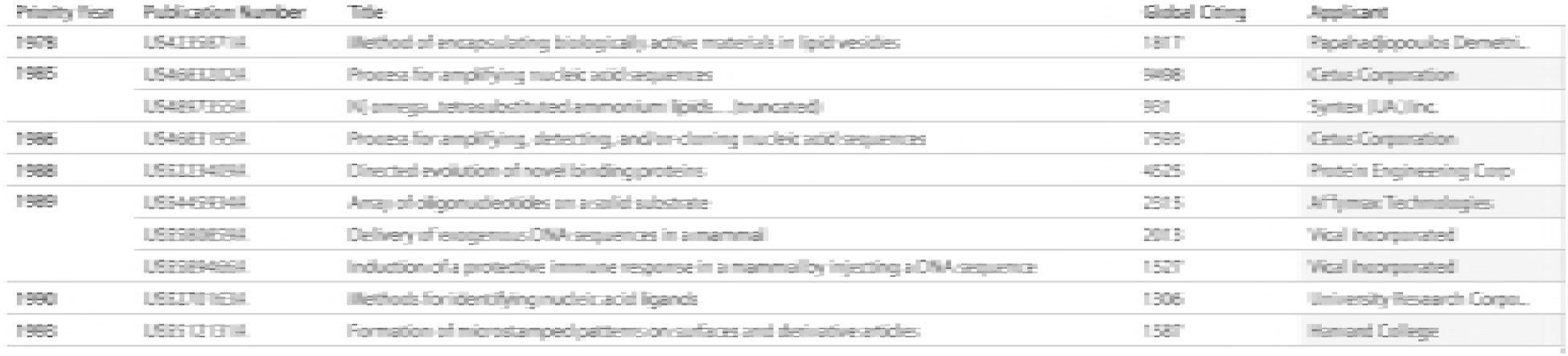
Top Cited Historic Patents in Synthetic Biology.

One of the challenges confronting researchers working on emerging technologies is where to begin. In the case of synthetic biology we could potentially begin with a report in the year 2000 declaring the début of synthetic biology [46]. As an alternative we might choose 2004 as the year of the first International Genetically Engineered Machine competition (iGEM). However, as Figure 3 reveals patent activity by researchers publishing on synthetic biology in the 21st Century can be traced to at least 1980. Furthermore, in approaching patent activity we need to recall that patent rights may endure for 20 years in many countries.

In interpreting the long tail of patent activity by researchers who have subsequently published in the field of synthetic biology it is useful to recall Wacław Szybalski’s comments from a 1973 panel discussion at a conference on strategies for the control of gene expression.

> “Let me now comment on the question”what next”. Up to now we are working on the *descriptive* phase of molecular biology. We are unravelling and describing the viral control mechanisms as found in nature. But the real challenge will start when we enter the *synthetic* phase of research in our field. We will then devise new control elements and add these new modules to the existing genomes or build up wholly new genomes. This would be a field with an unlimited expansion potential and hardly any limitations to building”new better control circuits” and “new better lambdas”, or finally other organisms, like a “new better mouse” instead of a better mouse trap. I am not concerned that we will run out of exciting and novel ideas, first in the *descriptive* and then in the *synthetic* phase of lambdology.” ([47] at 405, original emphasis)

Szybalski’s comments usefully summarise the history of molecular biology and genetic engineering in terms of a phase shift as part of a longer term agenda. Szybalski also anticipates much of the language of synthetic biology in terms of control elements, circuits, modules and the building or engineering of new genomes. In practice, the transformations and developments involved in these phases may involve decades and be punctuated by enabling technologies reflected in the cited landscape such as PCR, the ability to work with oligonucleotides, advances in the speed and accuracy of whole genome sequencing, the movement towards *in vitro* genome synthesis, the modification or design of DNA analogs and construction of genetic circuits and pathways, among others. When viewed from this perspective synthetic biology may more accurately be approached as a phase shift within biotechnology and genetic engineering taking place over decades punctuated by a series of scientific and technical landmarks. The use of CRISPR/Cas9 as molecular scissors in genome editing is one of the more recent landmarks. As we will see these developments are captured in the core landscape.

### Technology Areas and Topics

As noted above the patent system employs classification systems to describe the contents of documents and patent documents typically receive more than one classification code. The main classification is the International Patent Classification (IPC). Figure 7 displays the main technology areas for the core dataset at the sub-class level. Counts are based on first filings to avoid over counting. As IPC descriptors are typically sentences we have shortened the descriptions for ease of presentation.

**Figure 7:**
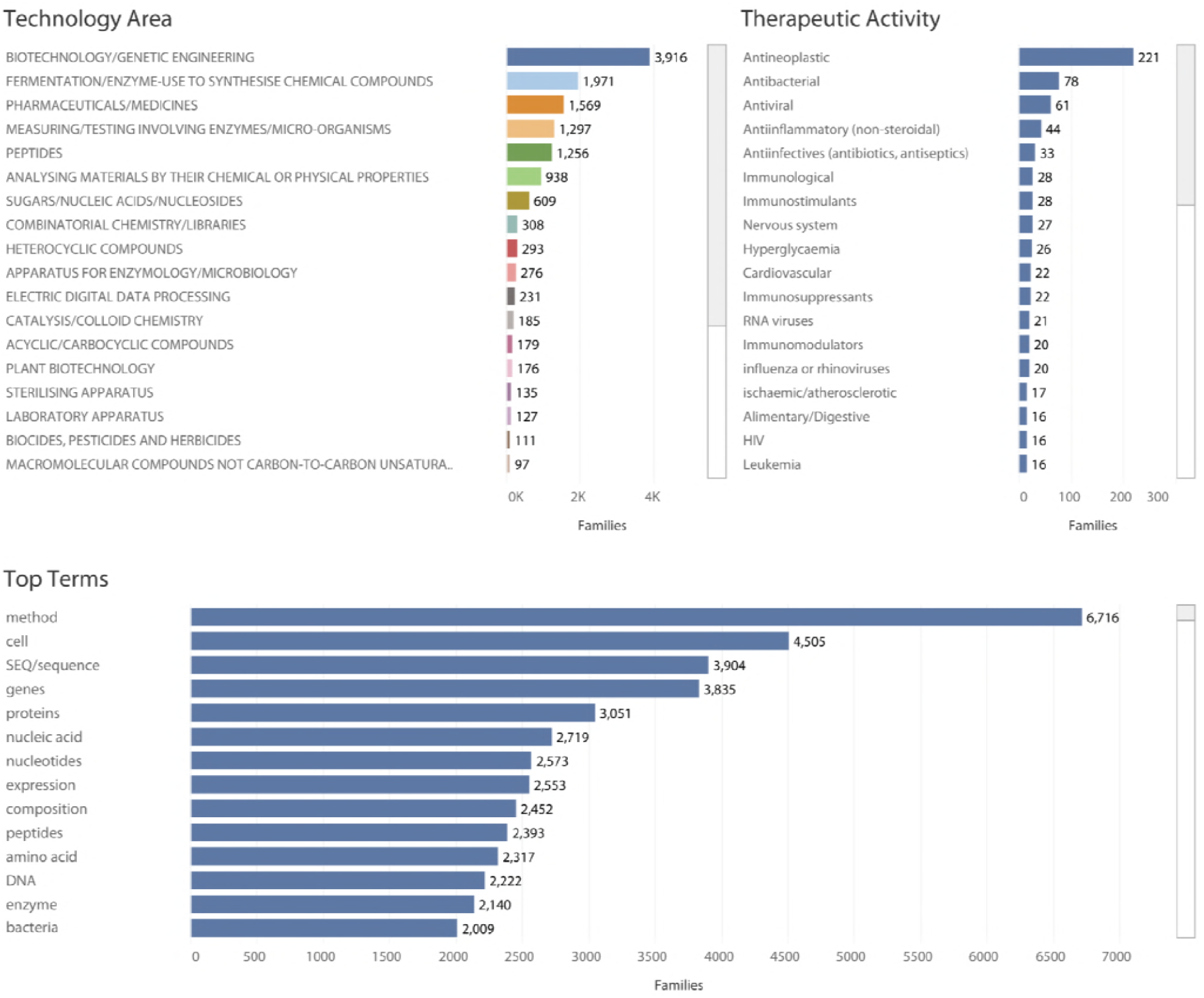
Top Technology Areas by IPC Subclass.

We can clearly see that biotechnology and genetic engineering is the main sub class (C12N) [26]. This is consistent with the view that synthetic biology is not readily distinguishable from biotechnology and an understanding of synthetic biology as the synthetic phase of molecular biology. This is also reflected in the prevalence of fermentation and the use of enzymes to synthesise compounds (C12P), along with measuring and testing involving enzymes or microorganisms (C12Q, the main classifier for DNA sequencing), peptides (C07K), analysing materials by identifying their chemical or physical properties (G01N), sugars, nucleic acids and nucleosides (C07H) and combinatorial chemistry (C40B). As we can also observe pharmaceutical and medical applications (A61K) dominate applications followed by plant biotechnology (A01H), biocides and pesticides (A01N), diagnostic tools (A61B), animal husbandry (breeding) (A01K) and foodstuffs (A23L). As this suggests, at the level of classification, synthetic biology is firmly situated within the biotechnology space with important connections in areas such as nanotechnology (B82B, B82Y) and computing (GO6F).

At the level of classification codes, coverage of specific uses of claimed inventions is limited. In the case of medical applications examiners may use descriptive codes (A61P) to describe the therapeutic activity. While these codes are not awarded to all documents with medical applications Figure 7 provides an insight into the orientation of research focusing on pharmaceuticals and medicines.

We can gain a more detailed view by text mining the documents to identify subjects of interest. Text mining can be performed across the entire text or the title and abstract or title, abstract and claims. Text mining of the full text is beyond the scope of this article. However, a common working practice in patent analytics is to capture the universe of full text documents in the first instance and then focus on the title, abstract and claims to identify the main subject matter in a refined way.

We reviewed 606,794 words and phrases from the titles, abstracts and claims with a specific focus on all terms appearing in 20 or more records. Drawing on our earlier work the data was aggregated on top frequency terms and those relating to uses and classes of organism (human, animal, plant, eukaryote etc.). This approach is intended to identify subjects of interest to the synthetic biology and wider research community on a broad level to assist with further investigation. A terms table is made available with the public dataset for joining to the core set to assist with filtering the data and methodological development. Our focus here is identifying some of the features revealed by text mining.

Figure 7 displays the top ranking terms appearing in the titles, abstracts or claims of the core landscape. Figure 8 displays the dense network of connections between the terms in the titles, abstracts and claims.

**Figure 8:**
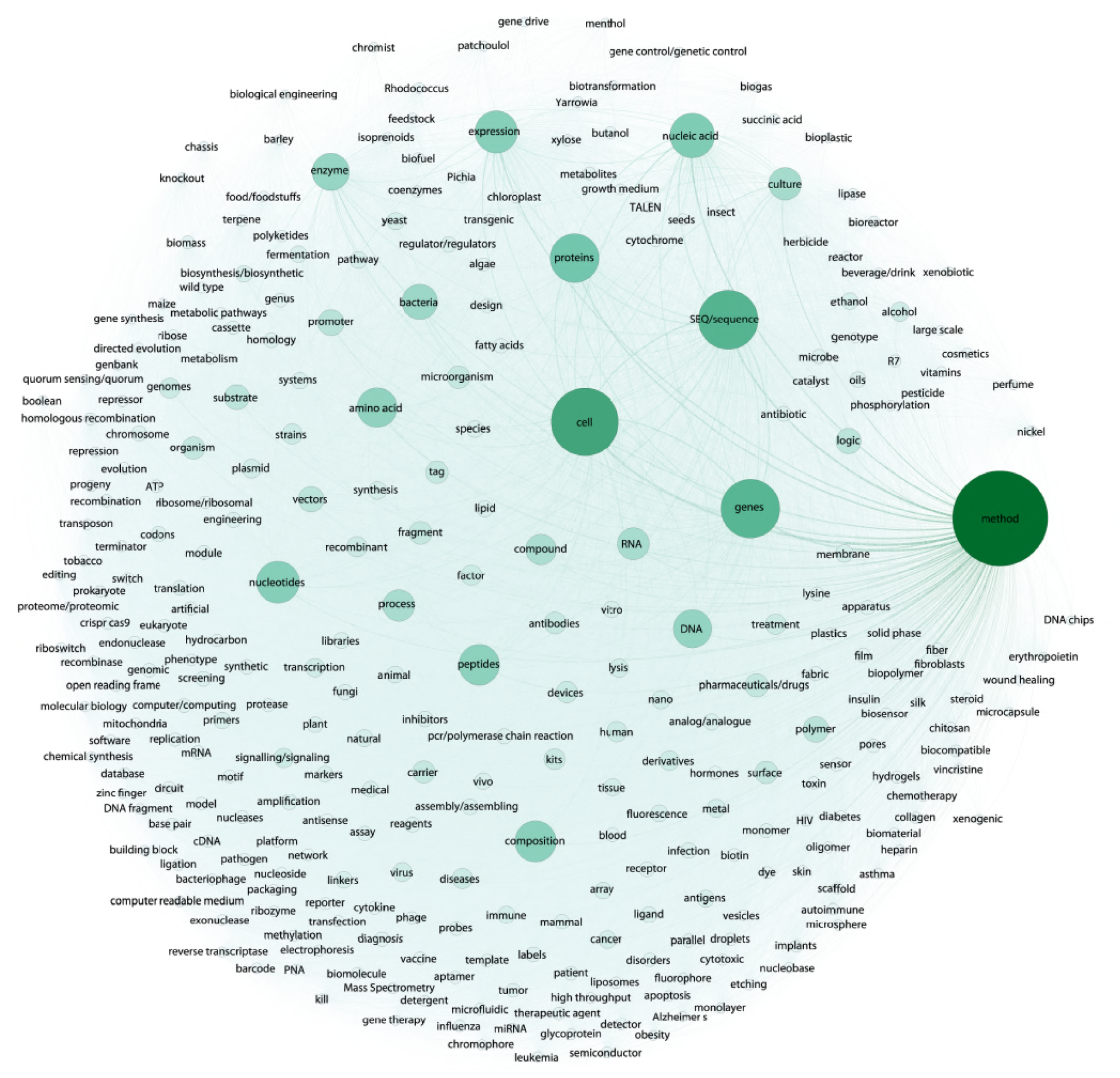
Top Terms in the Titles, Abstracts or Claims of the Core Landscape.

Figure 8 reveals that dominant terms in the titles, abstracts and claims aggregate around the word method along with cells, SEQ/sequence, genes, proteins, nucleic acids, nucleotides, expression (e.g. gene expression), compositions (for compositions of matter), and peptides.

The prominence of terms such as method, composition and process are directly linked to the three main types of patent claims for: methods, compositions of matter, and processes (products by process). For example:

> “A computer-implemented method for identifying a regulatory interaction between a transcription factor and a gene target of said transcription factor, the method comprising…” (US8306752B1)

In raw terms references to compositions appear second. However, in practice the majority of claims (approximately 4,359) are constructed as compositions of matter and begin with words such as “A cell”, “a compound” etc. An example of a composition of matter claim is as follows:

> “A non-naturally occurring or engineered self-inactivating CRISPR-Cas composition comprising; a first regulatory element operably linked to a CRISPR-Cas system RNA polynucleotide sequence…” (US20160354487A1)

Process focused applications are much less frequent in the data and often associated with methods. An example of a process focused patent application involves carbon capture with chemoautotrophic organisms:

> “A biological and chemical process for the capture and conversion of carbon dioxide and/or other sources of inorganic carbon, into organic compounds, comprising: introducing carbon dioxide gas, either alone and/or dissolved in a mixture or solution further comprising carbonate ion and/or bicarbonate ion, and/or introducing inorganic carbon contained in a solid phase into an environment suitable for maintaining chemoautotrophic organisms and/or chemoautotroph cell extracts…“ (US20130078690A1)

The prominence of these terms relates to how patent claims are constructed. In addition, we observe a high frequency of the terms SEQ/sequence in the claims (approximately 4067). These references are associated with claims to DNA and amino acid sequences. However, as Jefferson et. al. 2013 have highlighted, sequences may be mentioned in the claims either because they are being claimed or because they are reference sequences (not claimed) for the invention [48]. As such, while references to sequences are extremely common in patent claims more detailed investigation will be required to assess the significance of these references at the level of patent rights. As these observations highlight, debates on the implications of patent activity for synthetic biology will require careful consideration both of the types of claims that are being made and the legal significance of references to sequences in patent claims.

High frequency terms within the core landscape reinforce the point that synthetic biology is probably best understood in terms of a phase shift within biotechnology and genetic engineering. However, less frequent terms such as those associated with genetic circuits, biosensors, biosynthetic or metabolic pathways, directed evolution, CRISPR CAS9, gene drives and DNA analogs speak more directly to distinctive areas of research. Figure 9 displays a selection of records in connection with genetic circuits, genome engineering and gene drives. In the case of gene drives, the use of this term is presently limited with other terms relating to expression systems, insects and germlines required to expand data capture. We will highlight some of these features in more detail in the exploration of inventor networks below.

**Figure 9:**
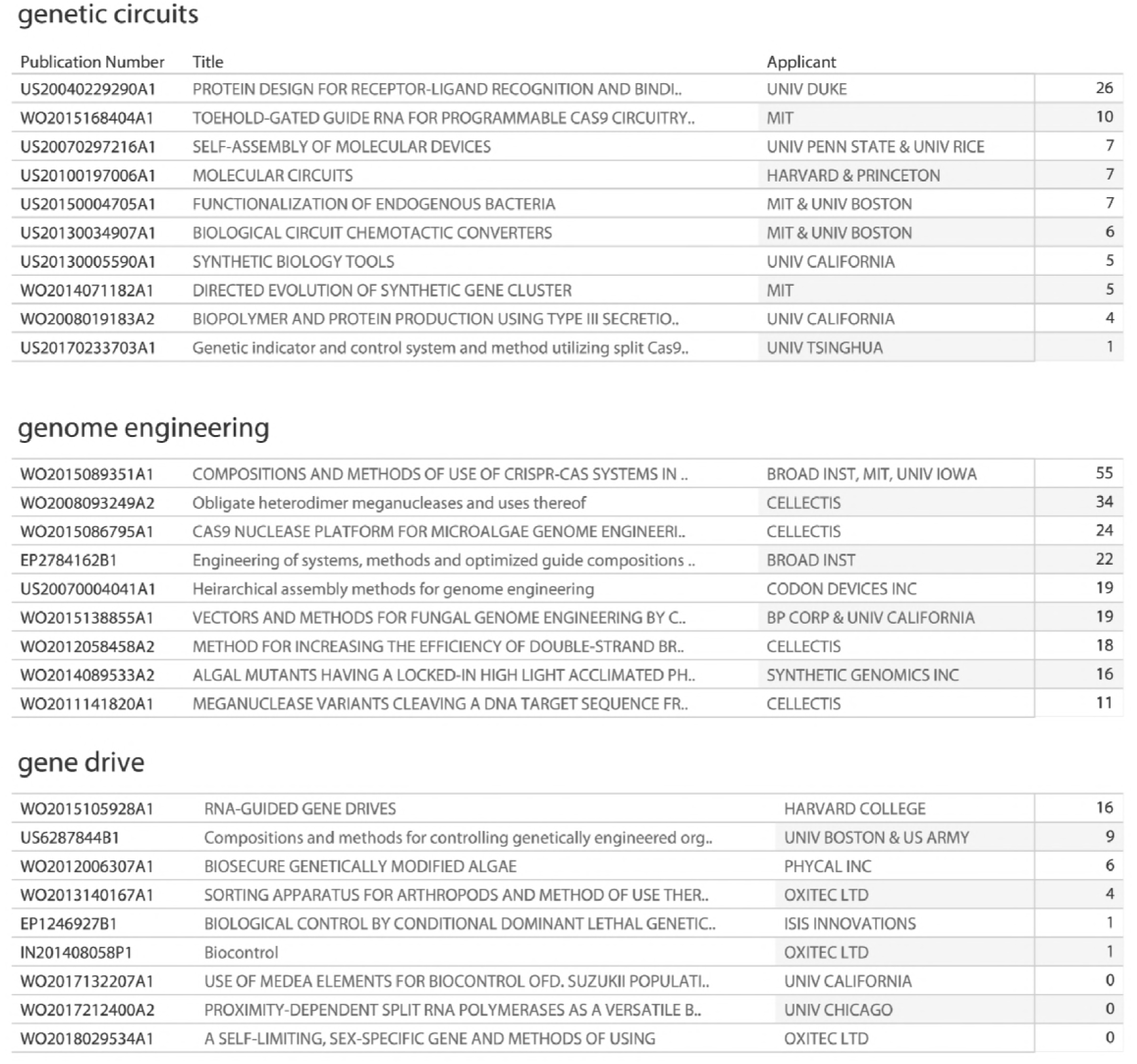
Examples of Selected Themes.

When approaching new and emerging areas of science and technology it is easy to gain the impression that patent activity is increasing rapidly. However, understanding patent trends depends on the type of measure that is used. In the next sections we begin by examining trends in unique first filings (families) of new patent applications and then turn to analysis of how those patent applications generate multiple other applications and grants worldwide.

### Trends in First Filings

The first filings of patent applications that form the basis for patent families are widely used in patent statistics as a proxy indicator for investments in research and development [49]. That is, the first filing of a patent application is an output indicator for investments in research and development that may otherwise be invisible [49]. As we will see, a degree of caution is required in the interpretation of this data on a national level due to the variety of routes that applicants may use when filing patent applications and problems of timeliness in the availability of patent data.

Figure 10 displays the origins of the first filings of patent families by country across all years for the core landscape. Figure 11 displays trends in first filings for the top ranking countries and instruments. In considering Figure 11 note that the y axis for each country is independent.

**Figure 10:**
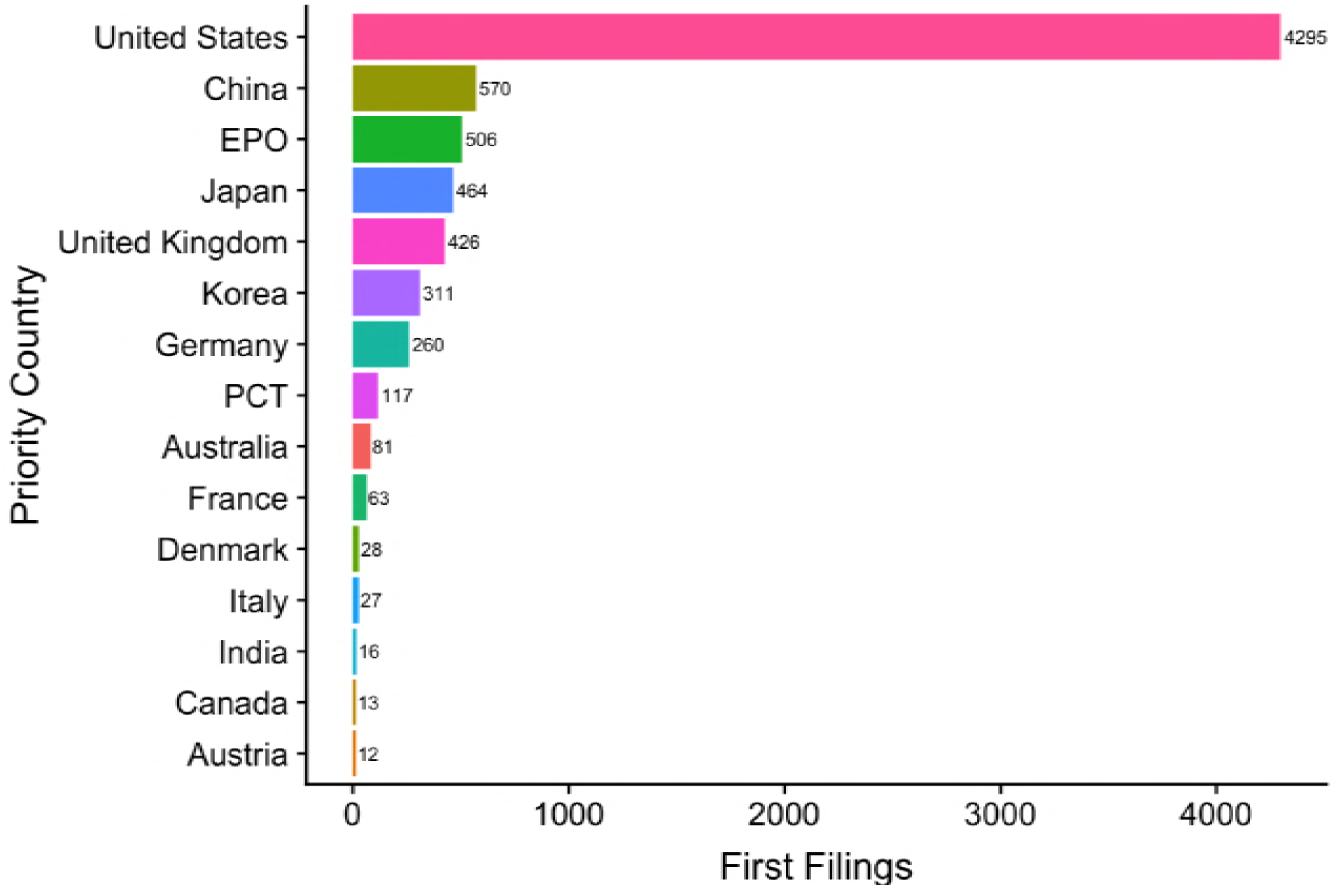
First Filings by Country.

**Figure 11:**
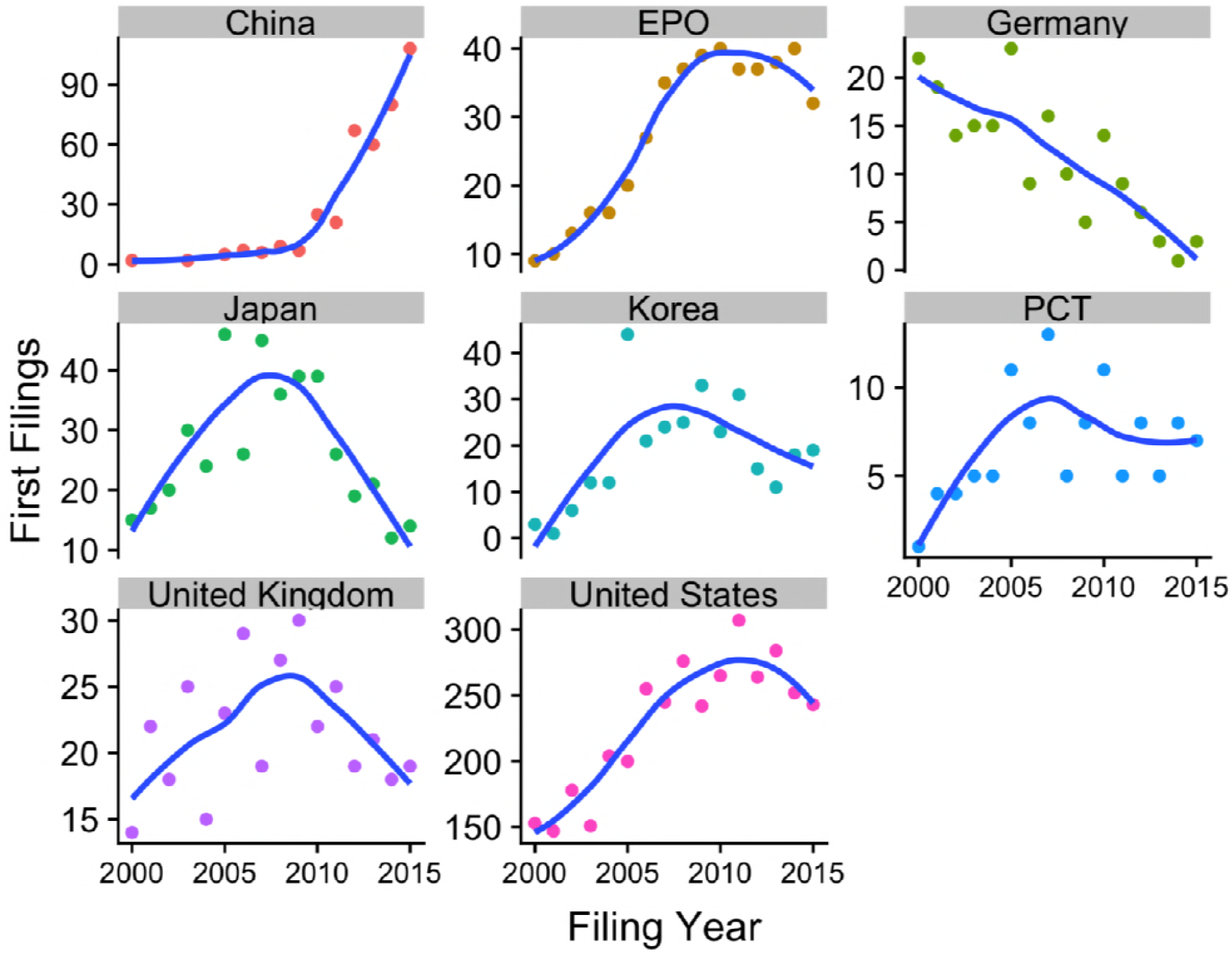
Trends in First Filings by Country.

Figure 10 reveals that the United States and China rank top in terms of the number of first filings across the core landscape, followed by filings under the regional European Patent Convention (EP), Japan and the UK. Figure 11 plots trends in the first filings of patent applications across the top countries using the earliest filing data for the period 2000 to 2015.

A notable feature of Figure 11 is that while trends in first filings are rising steeply in China (from a low base) they appear to be levelling off in the United States. In other cases we observe declining trends in national level filings originating within a country, notably in the case of Germany, the UK, Japan and Korea. However, we would caution that national level priority filings may not reveal the whole picture for two main reasons. First, applicants can increasingly choose different filing routes. In the case of Germany a total of 260 first filings were recorded in Germany. However, applicants and inventors from Germany also appeared on 239 first filings of regional applications at the European Patent Office (which has offices in Germany). As such, applicants are increasingly pursuing the regional route that will not appear in national counts of first filings. In contrast applicants from the UK appear on only 28 European filings suggesting that filings relating to synthetic biology in the UK are presently limited. However, as we will see below, the declining trend in first filings for Germany is also mirrored in a declining trend at the level of patent family members and reflects a wider decline in patent activity for biochemistry in Germany in recent years. The second reason for caution relates to the problem of timeliness. That is, the closer we move to the present the less data is available. In the case of countries such as the UK significant investments in synthetic biology programmes are relatively new and it may be that these investments are not yet reflected in the patent data as an output indicator. While the data is correct, these reservations expose some of the underlying issues involved in developing accurate national level patent statistics and suggests a need for the development of national level studies to improve the resolution of the picture.

Figure 11 addresses the data on trends in filings originating within a country or under a patent instrument based on the first filings. As we have seen, and bearing in mind that the majority of researchers in synthetic biology do not file for patent protection, trends in filings from within a country may actually go down over time. However, this is only part of the national level picture. Because the patent system is global in nature, applicants may pursue protection in multiple countries. As a result, national level trends and portfolios for synthetic biology may in reality be dominated by demand from third countries rather than filings by nationals. We now turn from trends in filings to the analysis of trends in demand for protection.

### Global Trends in Demand for Patent Rights

First filings of patent applications that form the basis of patent families are typically published as applications two to three years after they are filed. They may also then be republished as patent grants or as administrative republications (corrections, search reports etc.). As we have noted above patent applicants increasingly make use of regional and international patent instruments, notably the European Patent Convention and Patent Cooperation Treaty. These instruments allow applicants to submit a single application that may potentially be considered in multiple other countries. In the case of the Patent Cooperation Treaty a single patent application could potentially go forward for consideration in up to 152 Contracting States. However, there is no ‘global’ patent and each time that an applicant decides to pursue protection in a specific country an application will normally be republished and, if successful, republished as a patent grant. This introduces radical multiplier effects into patent counts but also allows us to understand global trends in demand for patent rights. The republication of patent documents is an indicator of demand for patent rights because applicants will normally pay fees at each stage of the procedure in countries where they seek protection. As such, the number of patent family members is an expression of willingness to pay for protection.

The relationship between first filings and global demand for protection can be mapped using patent family relationships. The first filings in the core landscape are grouped into 7,424 patent families that are linked to 71,887 family members worldwide [44]. Figure 12 displays global demand for patent rights across all years as reflected in the number of family members linked to the first filings.

**Figure 12:**
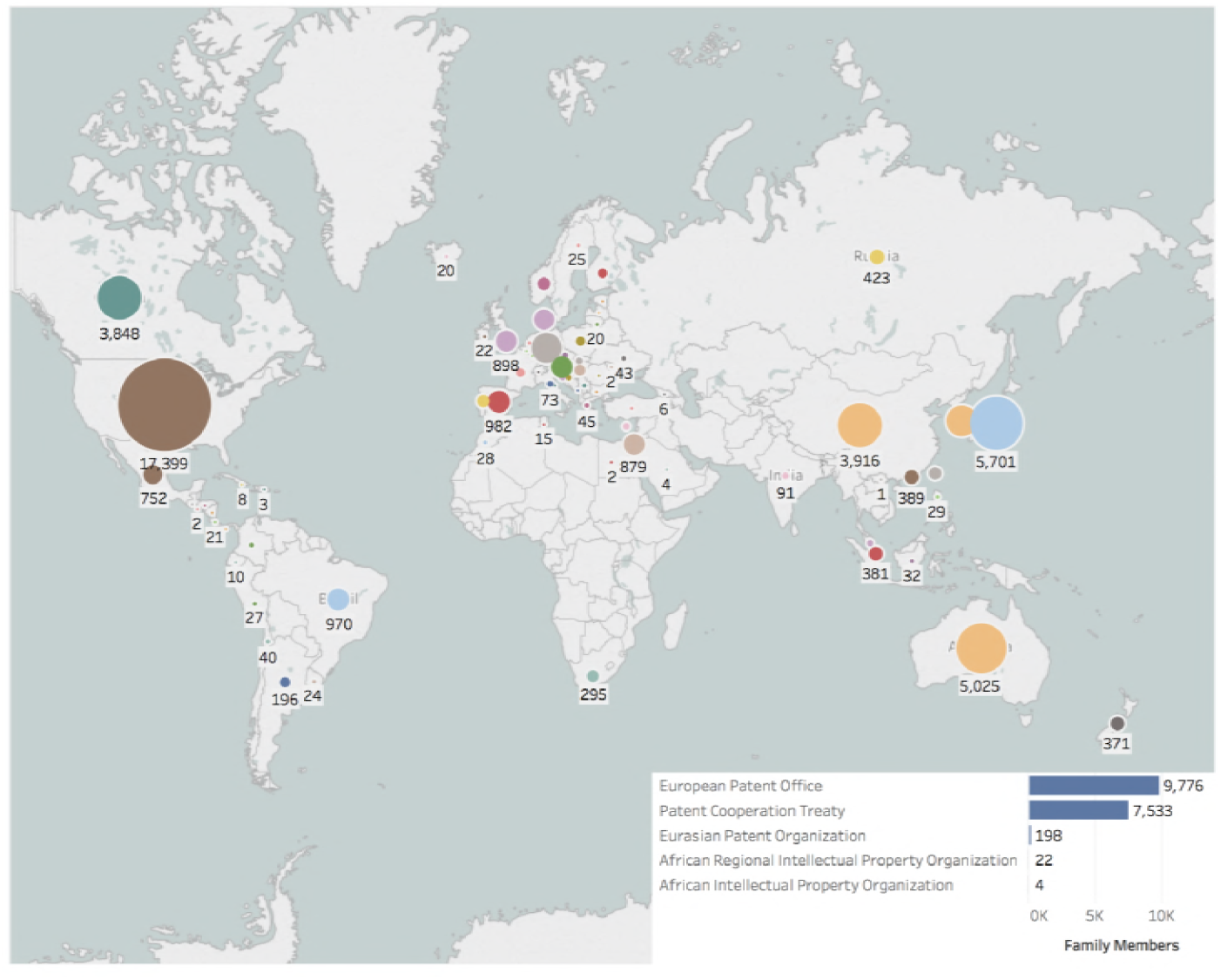
International Demand for Patent Rights.

In considering Figure 12 we observe, as expected, that demand is concentrated in the United States followed by Japan, Australia, China and Canada. Our view of data from countries such as India and South East Asia is limited due to the lack of visibility of national collections from these countries to the INPADOC family system. Figure 13 displays this data organised by region and income level using the World Bank Development Indicators classification [50].

**Figure 13:**
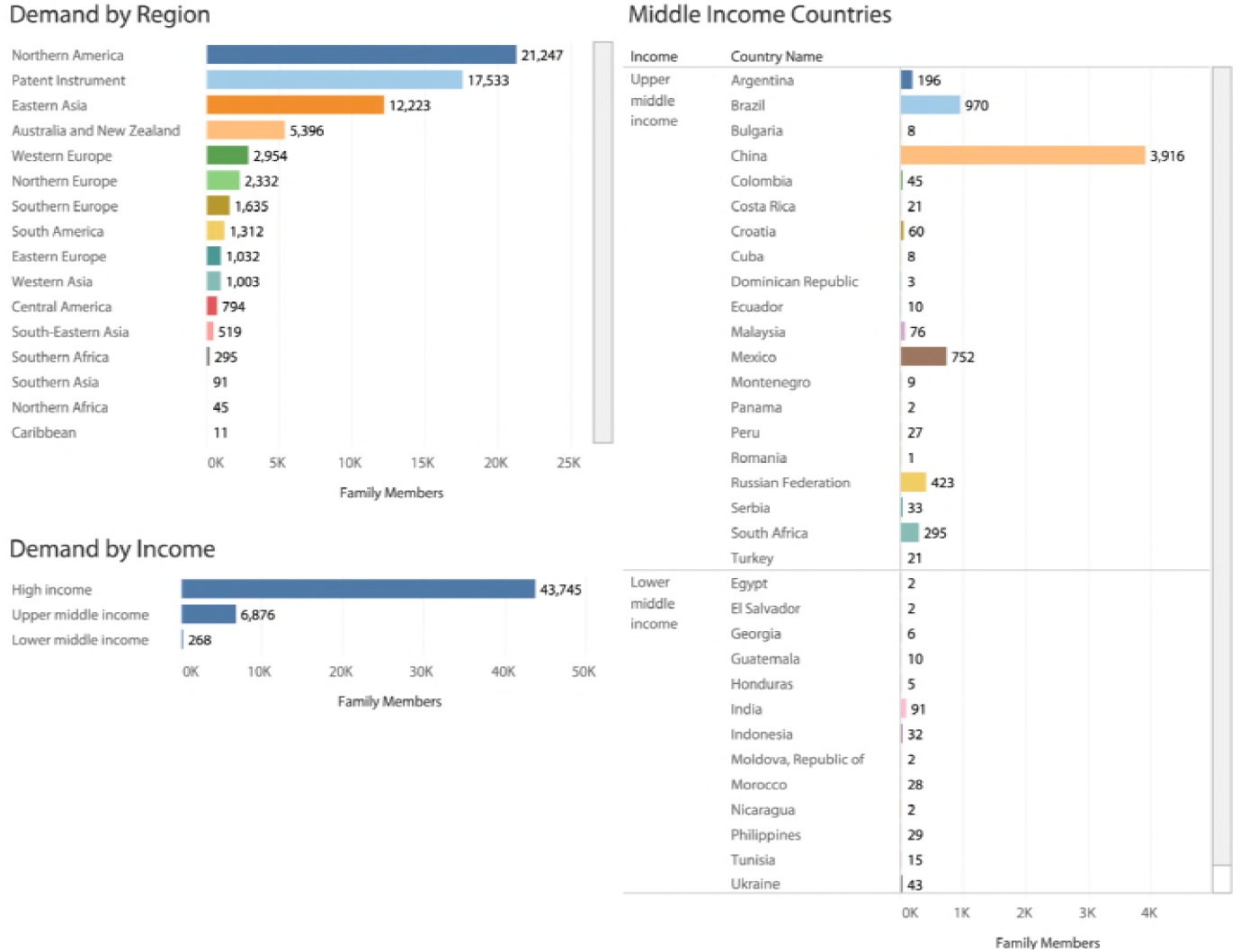
Demand for Patent Rights through Patent Family Members.

Figure 13 demonstrates that demand for protection across the core dataset, within the limitations noted above, is concentrated in North America and East Asia, and in high income countries. In the case of middle income countries Brazil, Russia, China and South Africa (BRICS) are an emerging focus of demand while we have a partial view for India along with other countries classified by the World Bank as Lower Middle Income.

Our ability to view trends in patent applications and grants is affected by variations in how patent offices record patent publications using kind codes (e.g. A1, B1 etc) and requires close attention to national level practices. Figure 14 displays aggregated data on publications of applications and grants for the top countries. This figure should be viewed in combination with Figure 11 on first filings discussed above.

**Figure 14:**
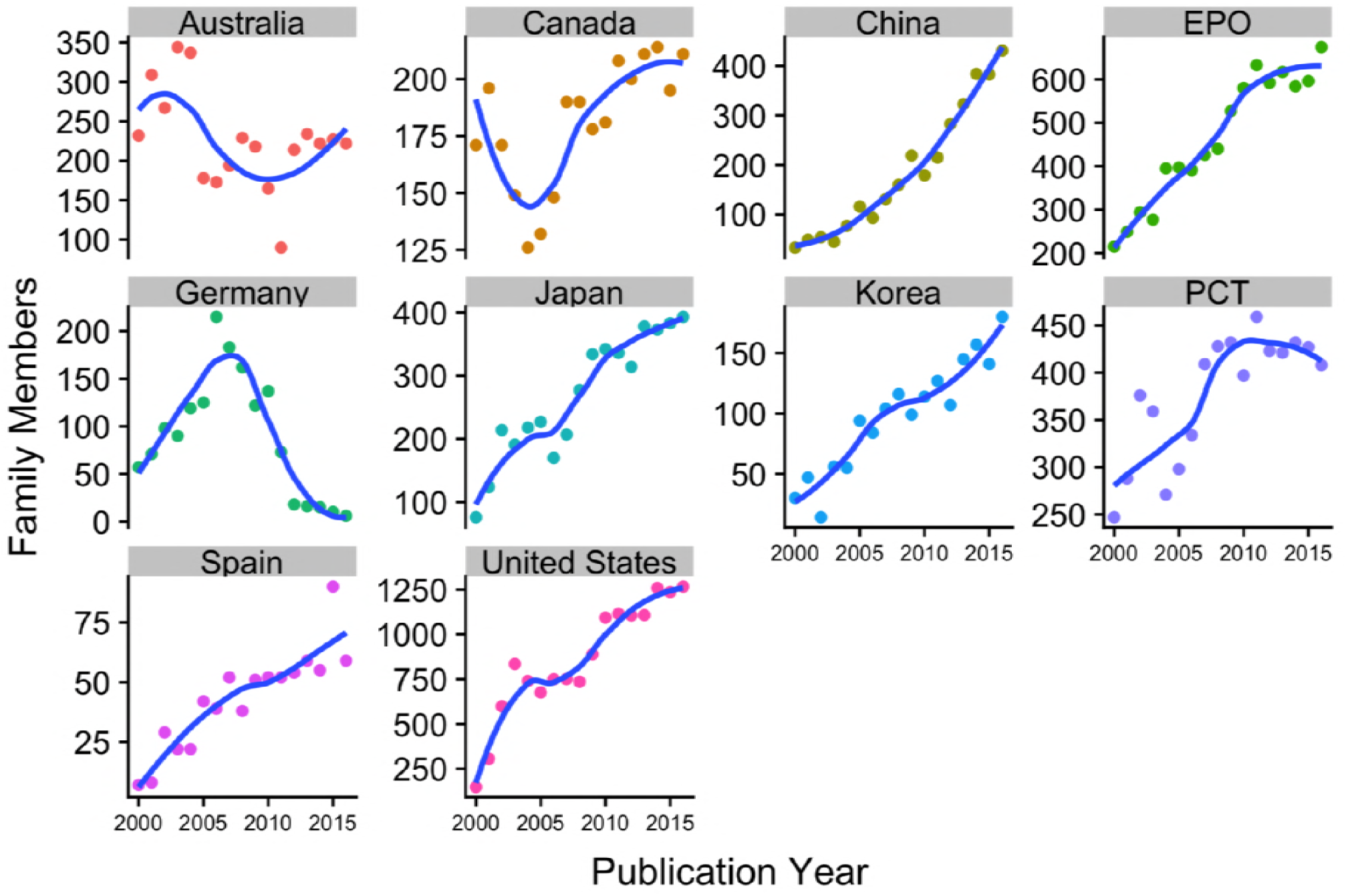
Trends in Demand for Patent Rights.

Note that the number of family members on the y axis in Figure 14 is significantly higher than filings in Figure 11. In the case of Australia we observe a significant trough in activity in the mid 2000s before a rising trend. This is likely to at least partly reflect issues reported in the patent analytics community involving the treatment of PCT applications in Australia in the opening years of the 21st Century (which counted PCT designations or potential applications as actual applications). In the case of Canada, the trough replicates a trough in overall publications in class C12 for Biochemistry in this period. The sharp declining trend in Germany also replicates the wider pattern for trends in Biochemistry (C12).

The remaining countries and instruments (with the exception of the Patent Cooperation Treaty) display steep rising trends in national level publications. When considered in light of national level first filings in Figure 11 this demand is primarily driven by third country applicants, notably from the United States, seeking protection in these markets. The main vehicle for this demand is the Patent Cooperation Treaty where a single application may lead to applications in multiple other countries.

The ability to identify national level trends is constrained by the complexities of national kind code systems. However, it is possible to map trends in applications and grants in the United States and Europe. Figure 15A displays trends in applications and grants in the United States while Figure 15B displays trends under the regional European Patent Convention.

**Figure 15:**
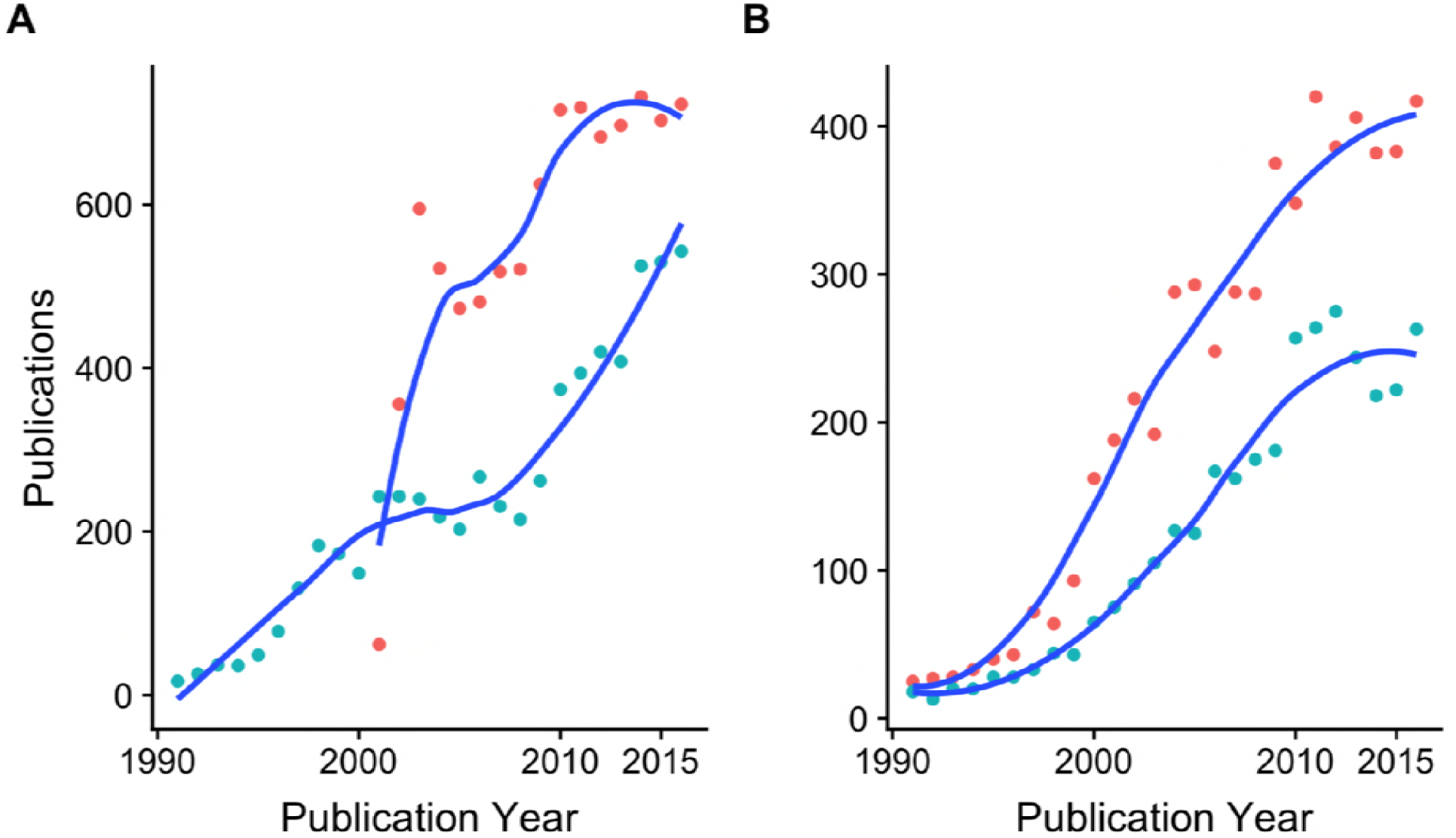
Trends in Patent Applications and Grants in the United States (A) and at the European Patent Office (B)

In considering the data for the United States two points should be noted. First, the USPTO only began publishing patent applications from 2001 onwards. Prior to this only granted patents were published. For this reason US publication data always displays a sharp spike from 2001. This is a reporting effect rather than an increase in actual activity. Second, US first filings are dominated by the use of the Provisional applications system [51]. Thus, 3699 of 4414 (83.8%) of US first filings are provisional applications. Under the provisional system applicants may submit an outline application with a description that does not include patent claims. Provisional applications are not published but establish a priority date relative to competitors. Provisional filings, and combinations of provisional filings, often form the basis for multiple other distinct applications. This is frequently combined with the practice of continuation, divisional and continuation in part applications [52,53]. Continuation applications commonly arise following objections from examiners and involve a new application based on the previous application with new claims. *Continuation in part* applications involve the addition of new subject matter and are commonly used to claim enhancements to the original claimed invention [54]. The use of continuation filings has the effect of repeatedly extending applications for essentially the same subject matter over time, inflating application data and arguably constitutes an abuse of the system [52]. Attempts by the USPTO to limit the number of continuation filings were abandoned in 2009 following litigation by opponents [55,56].

In the case of the European Patent Convention an applicant may submit regional applications covering the 12 member states and a regional level grant may be awarded. However, for a grant to have force member states commonly require national level translation of the grant and in effect, the result is a set of independent national level patent grants subject to national patent laws with a centralised regional level opposition system [57].

The analysis of trends in demand presented above reveals that demand is highly concentrated in developed countries with the exception of China and emerging activity in other BRICS countries. Within developed economies, demand for patent rights also varies with sharply increasing activity in some countries but a declining trend in countries such as Germany. The primary driver for patent activity is the international Patent Cooperation Treaty with the United States as the lead actor. A fuller picture is likely to emerge from country level studies. We now turn to analysis of patent applicants as a basis for in depth exploration of inventor networks.

### Patent Applicants

In total after name cleaning we identified 2,199 applicants, including individuals, across the patent landscape. Figure 16 displays the applicants ranked by the number of filings (families) with the second panel displaying applicants ranked by the number of family members.

**Figure 16:**
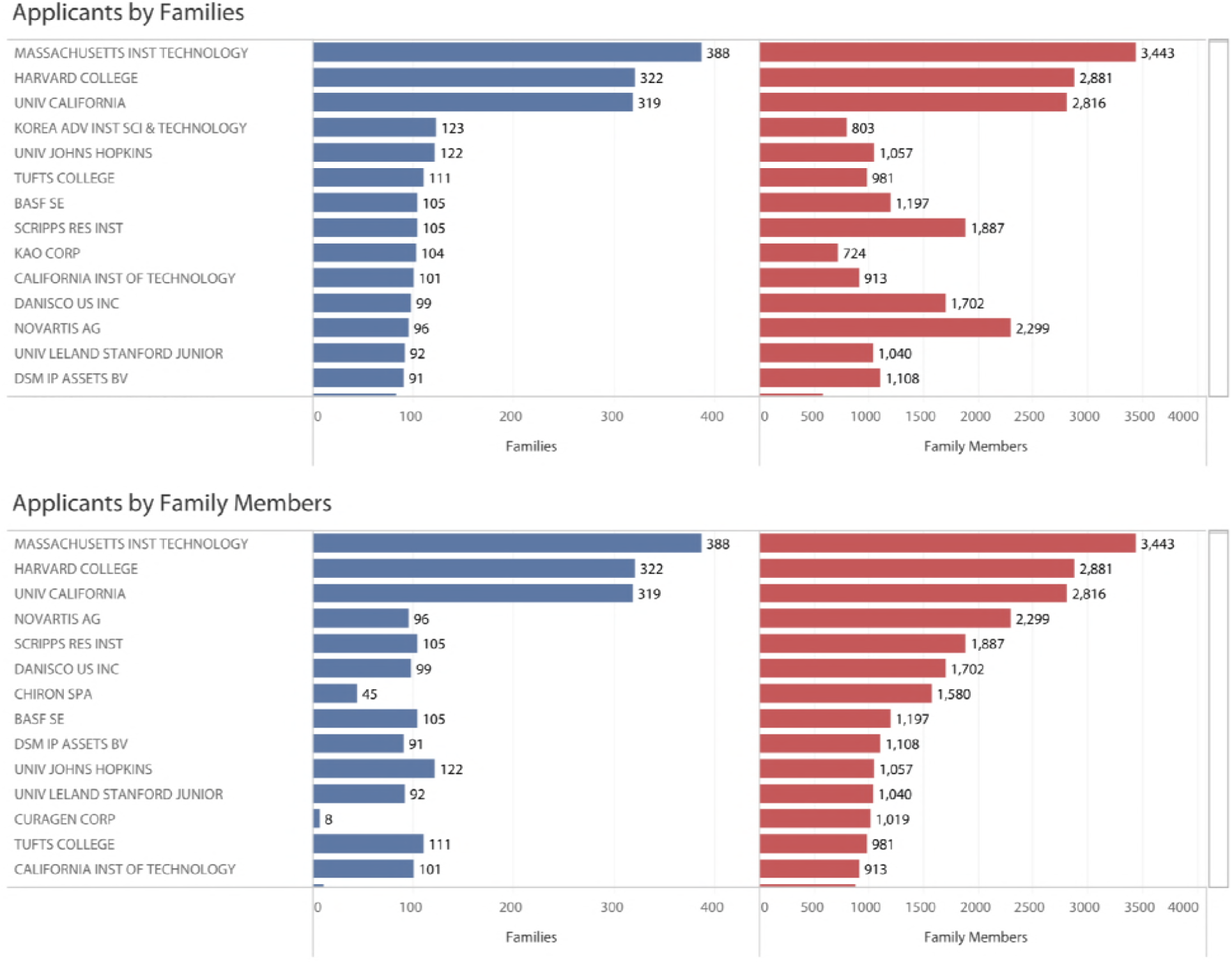
Patent Applicants.

Figure 16 reveals the dominance of MIT, Harvard and the University of California system in patent activity measured in terms of the number of first filings and reach in terms of the number of family members. However, when ranked on the number of family members, companies notably Novartis (see also Chiron which is now part of Novartis) become more prominent in the data and reflect the pursuit of patent protection in multiple jurisdictions. One company Curagen (acquired by Celldex in 2009) reveals a low number of first filings but a large number of family members, suggesting that this applicant intensively pursued protection for a limited number of inventions in connection with methods for analysing nucleic acids, sequencing of nucleic acids and proteins, with a focus on crop protection (CA2384510A1, CA2395341A1, US6388171B1, US6515202B1). Looking outside the United States, top applicants are revealed in Korea (Korea Advanced Institute for Science and Technology or KAIST), the German chemical company BASF, Kao Corp in Japan which focuses on chemicals and cosmetics, the US division of the Danish company Danisco (owned by DuPont) focusing on foods and ingredients, and the Dutch DSM that focuses on nutrition, health, personal care, medical devices and green products.

Patent applicants also form networks through joint applications for patent rights. These networks are typically sparse but assist with revealing collaborations between organisations. Figure 17 displays the network for organisations with five or more first filings (families). Node size is based on the number of families with the weight of edges reflecting joint patent applications. Nodes are coloured using a modularity class algorithm that identifies communities of closely related actors [58]. Note that due to the number of organisations in the network, edges may in some cases run through unconnected nodes.

**Figure 17:**
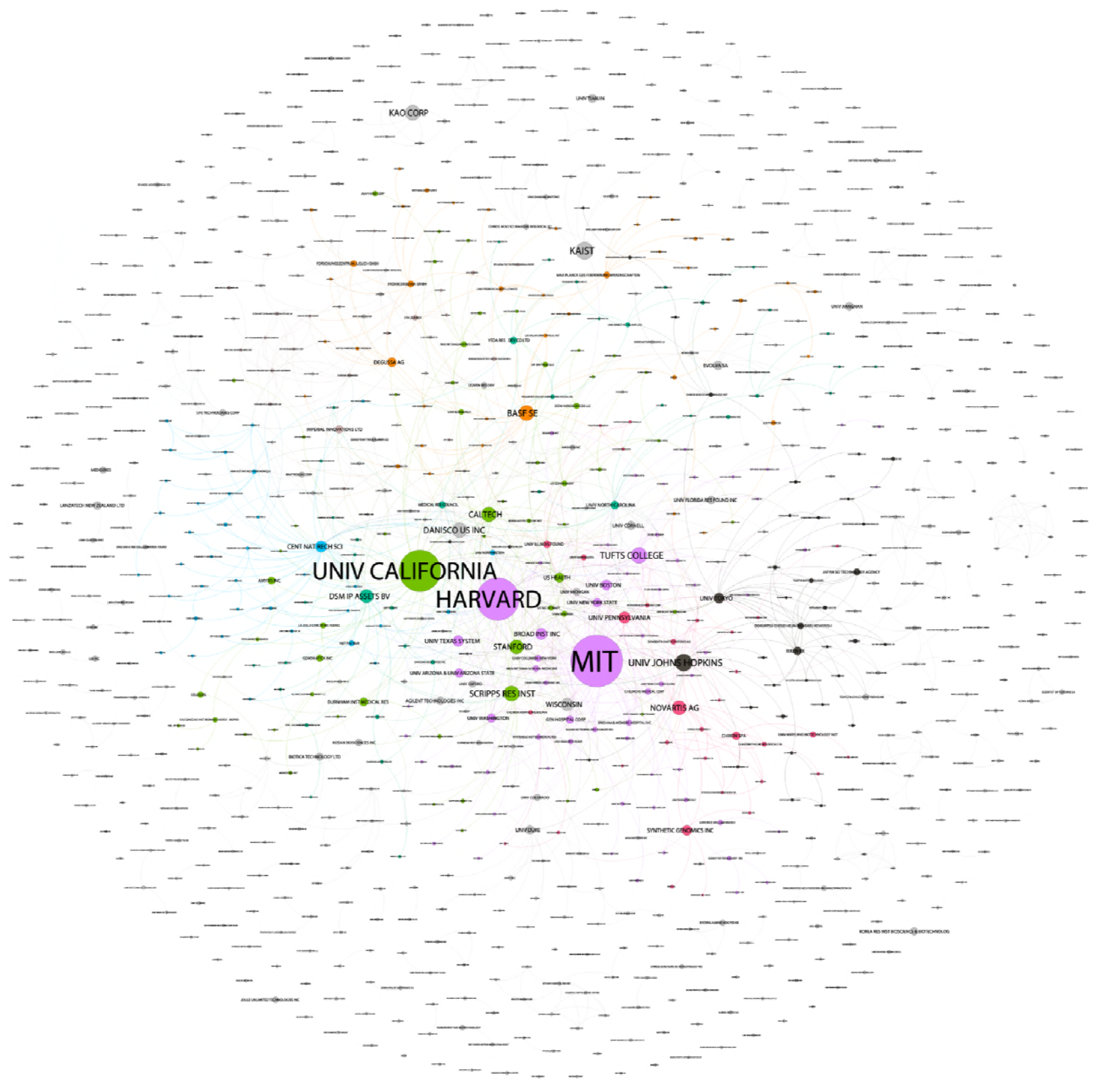
Patent Applicant Community Network.

Figure 17 reveals clear communities of joint applicants with core networks focusing on Harvard/MIT and the University of California system. Additional networks are also detectable such as Novartis, the J Craig Venter Institute and its commercial arm, Synthetic Genomics.

Figure 17 displays communities using colours. However, we can also explore the individual communities and the types of organisations involved in collaboration in research and development. To do this we colour organisations within the individual network by type: academic (green), company (purple), government (blue), hospital (orange). For ease of display the nodes are limited to those with four or more filings. In approaching these networks note that the division of organisations into four categories is imperfect because they may fall into more than one category.

Figure 18 displays the Harvard/MIT network. Figure 19 displays the University of California network and Figure 20 displays the Novartis/Venter Institute networks.

**Figure 18:**
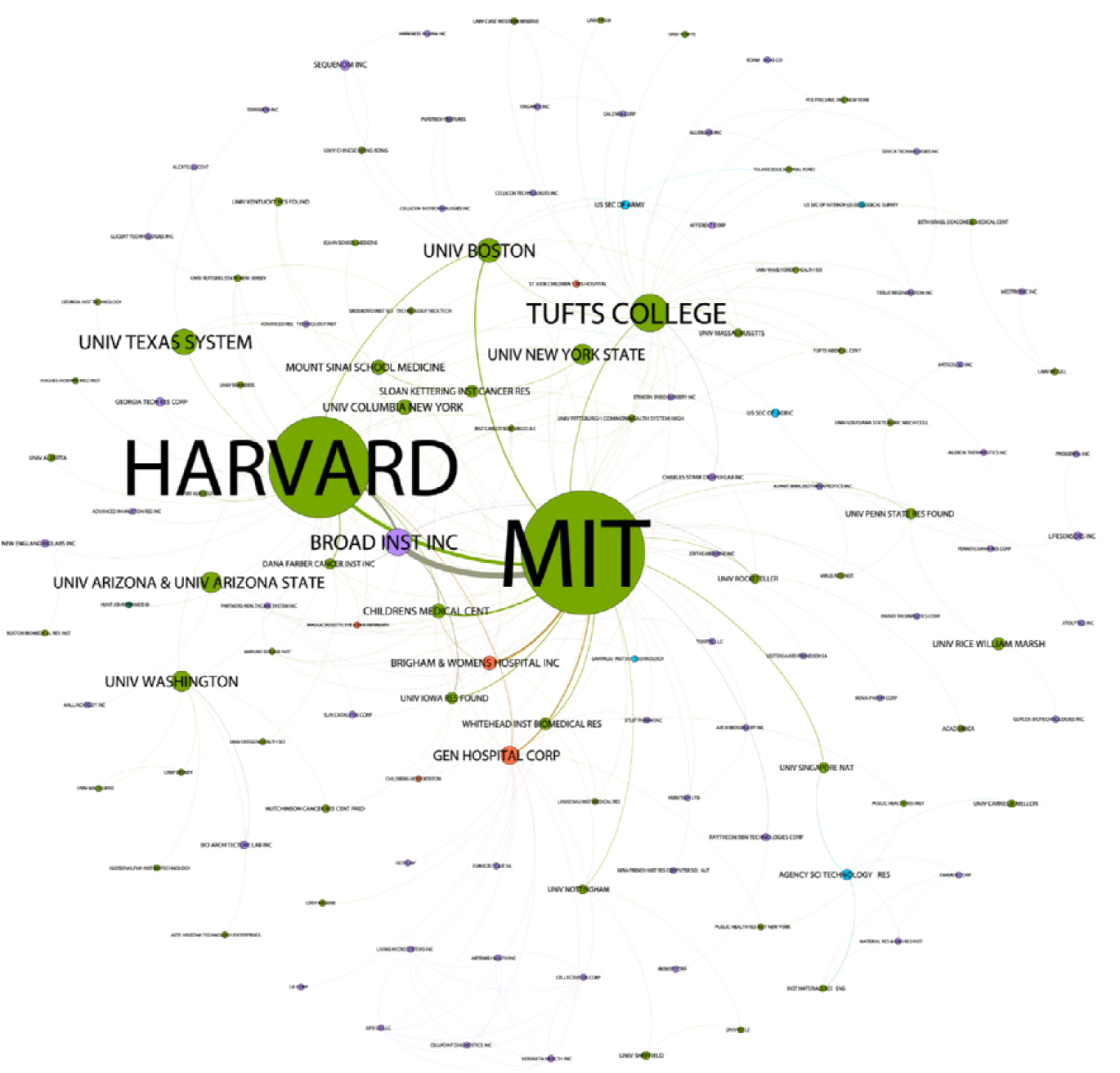
The Harvard/MIT Network by Organisation Type.

**Figure 19:**
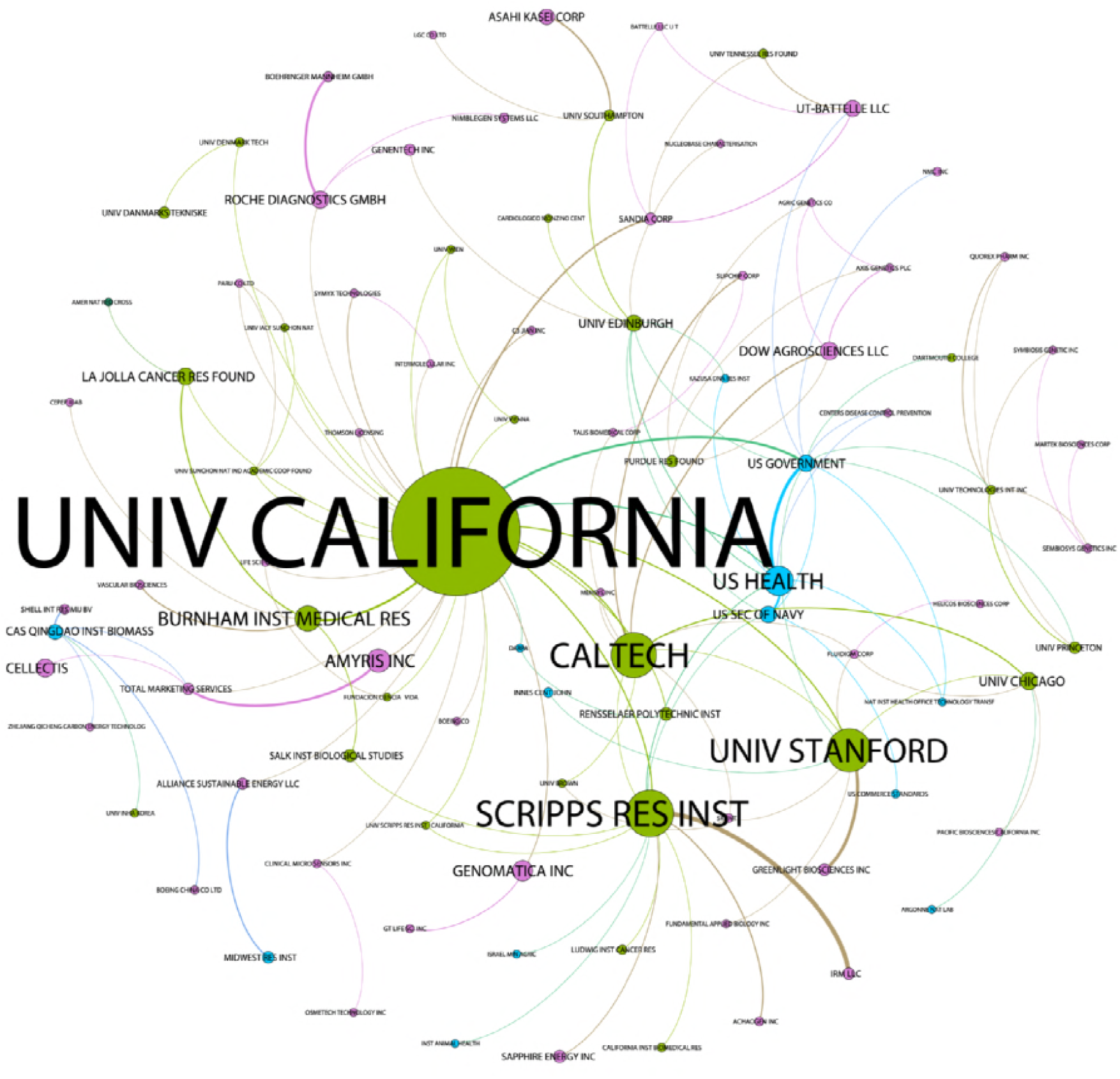
The University of California system network by Organisation Type.

**Figure 20:**
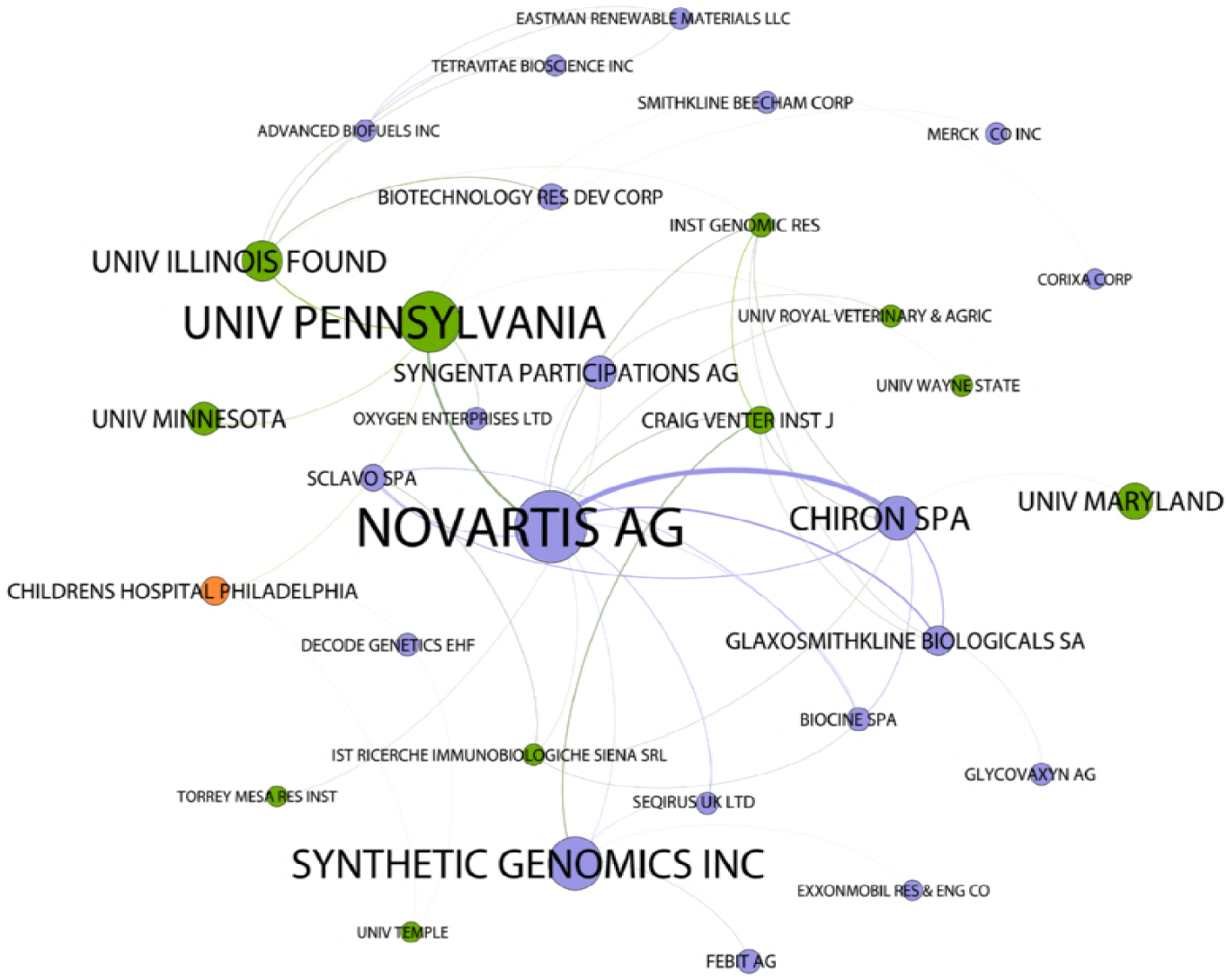
The Novartis/Venter Institute network by Organisation Type.

In approaching these networks it is important to emphasises that they are dynamic in nature. However, we can begin to explore the types of partnerships involved in the network. In the case of the Harvard/MIT network collaboration appears dominated by collaborations between academic institutions (in green), followed by companies (in purple). In practice, there is a strong orientation towards medical applications in the network that is somewhat underplayed by the classification scheme. Thus, in addition to Gen Hospital Corp, Brigham & Women’s Hospital, the Children’s Hospital Boston and the Massachusetts Eye & Ear Infirmary, we should add the Whitehead Institute for Biomedical Research (shown in Green) and the Broad Institute (shown in purple) to more clearly reveal a focus on health applications. These collaborations are clustered around the Boston area.

The University of California system network reveals a stronger collaboration with US government organisations in joint filings of patent applications when compared with the other networks. The network is also generally sparser and more concentrated among a smaller number of organisations in terms of joint patent applications than the Harvard MIT network.

In the case of the Novartis/Venter Institute network we observe the dominance of a commercial company (based on the number of filings) around which other actors such as the Venter Institute and Synthetic Genomics are articulated with linkages also observable to the Scripps Research Institute and University of Pennsylvania focusing on health applications. The presence of Syngenta suggests additional orientations towards agriculture and plant biotechnology while the presence of Sapphire Energy suggests a focus on collaborations in relation to biofuels (for the Scripps Institute).

What is striking in these network representations is that we are able to begin the process of identifying key partnerships in different sectors based on the core actors in these networks. For innovation studies this approach begins to expose empirical data on what is known as the triple helix of academic, industry, government relationships in innovation in synthetic biology [59,60]. Improvements in understanding these relationships in the case of synthetic biology could be achieved in future by focusing on the dynamics of network emergence and transformation over time.

Academic, Industry and Government patent collaboration networks are based on collaborations between researchers within and outside academia. Our approach is grounded in mapping authors of research on synthetic biology into the patent system. We now turn to analysis of patent activity by individual researchers and their collaboration networks.

### Inventor Networks

After name cleaning we identified 15,717 inventor names across the core landscape with 2,450 authors appearing as inventors. One challenge we encountered is that in some cases the most prominent inventors in the data, notably Robert Langer at MIT and Rino Rappuoli (formerly Novartis and now GSK), appeared to be peripheral in terms of the number of publications on synthetic biology in the baseline dataset. One approach to this issue would simply be to exclude authors below a threshold score (e.g. +1 publications). However, this ran the risk of potentially excluding important aspects of the landscape and relationships between actors. To address this issue we focused on approaching the data using five measures: a) counts of publications, b) counts of patent families, c) counts of patent family members as an indicator of global reach, d) counts of the number of co-authors in the literature dataset who appear as co-inventors, and; e) counts of citing patents as an indicator of impact on others within technology space. Figure 21 displays three ways of ranking authors of publications on synthetic biology who are also inventors.

**Figure 21:**
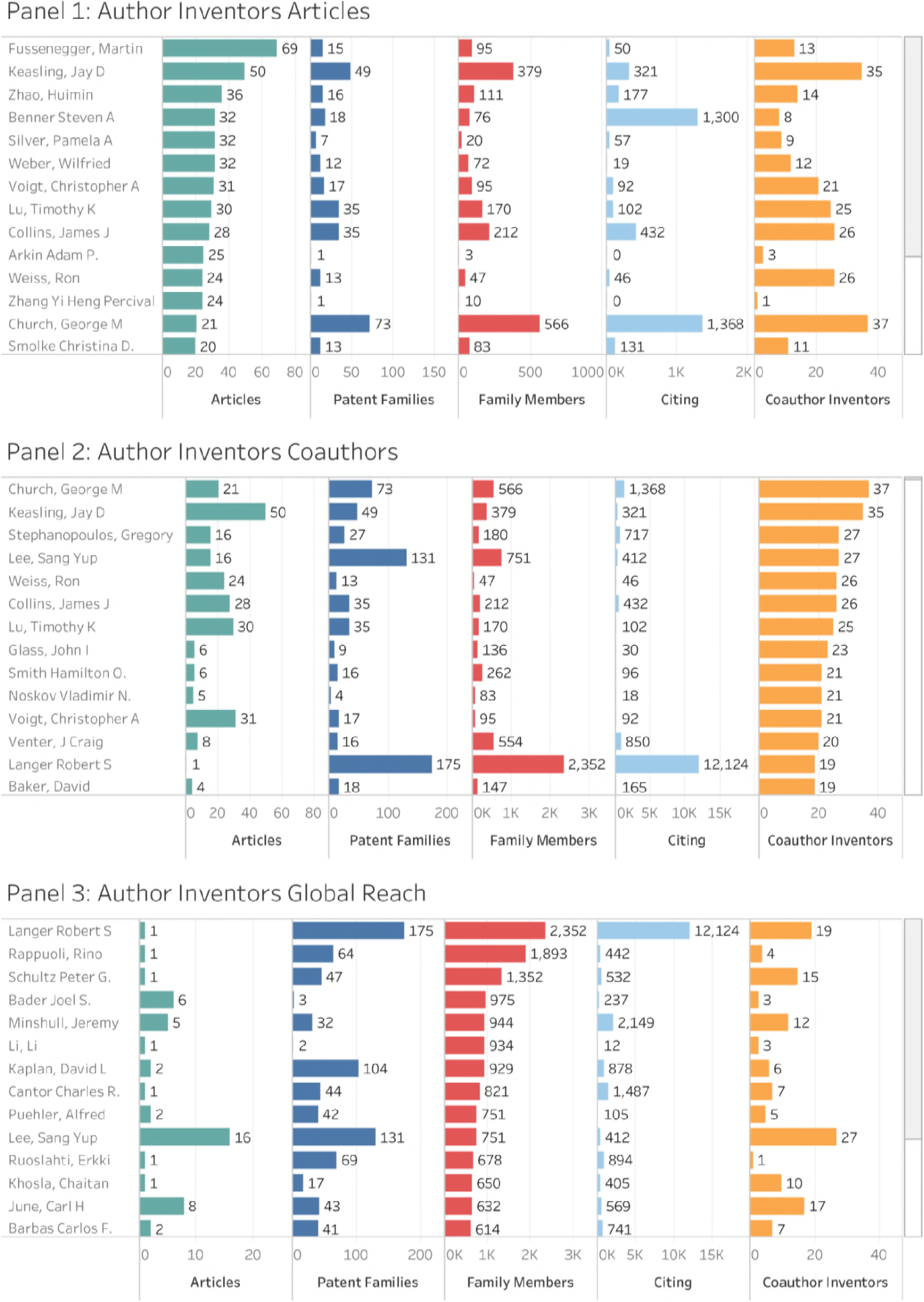
Author Inventor Rankings.

Panel 1 in Figure 21 displays authors who are inventors ranked on the number of articles that appear in the baseline literature dataset. This approach reveals that those authors who rank highly on the number of publications in the scientific literature in our baseline set are not necessarily highly ranked in terms of the number of patent filings. The most prominent of these is Martin Fussenegger who consistently appears as the top author in the baseline set but who displays limited patent filings in areas such as antibiotic based gene regulation (WO2000065080A1), SM protein based secretion engineering (WO2009080299A1), and a recent filing for a programmable differentiation control network in mammalian cells (EP3184638A1). Other author inventors in Panel 1 are also well known within the synthetic biology community and display considerable variation in the number of patent filings and global reach (family members).

Panel 2 in Figure 21 ranks author inventors based on counts of the number of coauthors across the dataset who appear as co-inventors within their respective patent portfolios. This measure focuses our attention on the intensity of collaboration between researcher groups in both publishing scientific research in a given field and pursuing its commercialisation through patent activity. For senior researchers, such as George Church at Harvard or Gregory Stephanopolous at MIT, this measure also perhaps serves as an indicator of the willingness of senior researchers to encourage younger researchers such as PhD students and post-docs to pursue commercialisation of research as part of their career development. Furthermore, we speculate that there is likely to be an association between coauthor clusters and the establishment of spin off companies. Rankings based on co-author inventors also bring members of the Venter Institute, notably John Glass, Hamilton Smith and Craig Venter to greater prominence.

Panel 3 in Figure 21 ranks author inventors by the number of raw patent family members. We present this as a measure of global reach because large numbers of family members will normally represent the pursuit of patent protection, typically by an employer or commercial partner, in multiple countries.

The most striking feature in Panel 3 is that researchers with low publication scores, notably Robert Langer at MIT, Rino Rappuoli, formerly at Novartis and now at GSK Vaccines, along with Peter Schultz at the Scripps Institute, dominate the landscape to a significant degree in terms of both the number of filings and family members in global reach. As noted above, when confronted with these at first surprising results it may be tempting to control results by the number of publications as a condition of entry into the dataset. However, as the co-author inventor scores reveal, the use of such simple control measures may miss the significant influence of senior researchers who may otherwise appear peripheral upon emerging fields. As we will see in further detail below, in the case of Robert Langer the network of author inventors is significant in areas such as genome editing while in the case of Rino Rappuoli the implementation of genomics based reverse vaccinology involved collaboration with members of the J Craig Venter Institute.

The foundation for our analysis is the identification of authors as inventors. The data in Figure 21 can also be visualised as a network. Once again we used a modularity class community detection algorithm to identify research clusters in the network displayed in Figure 22 using colours [58].

**Figure 22:**
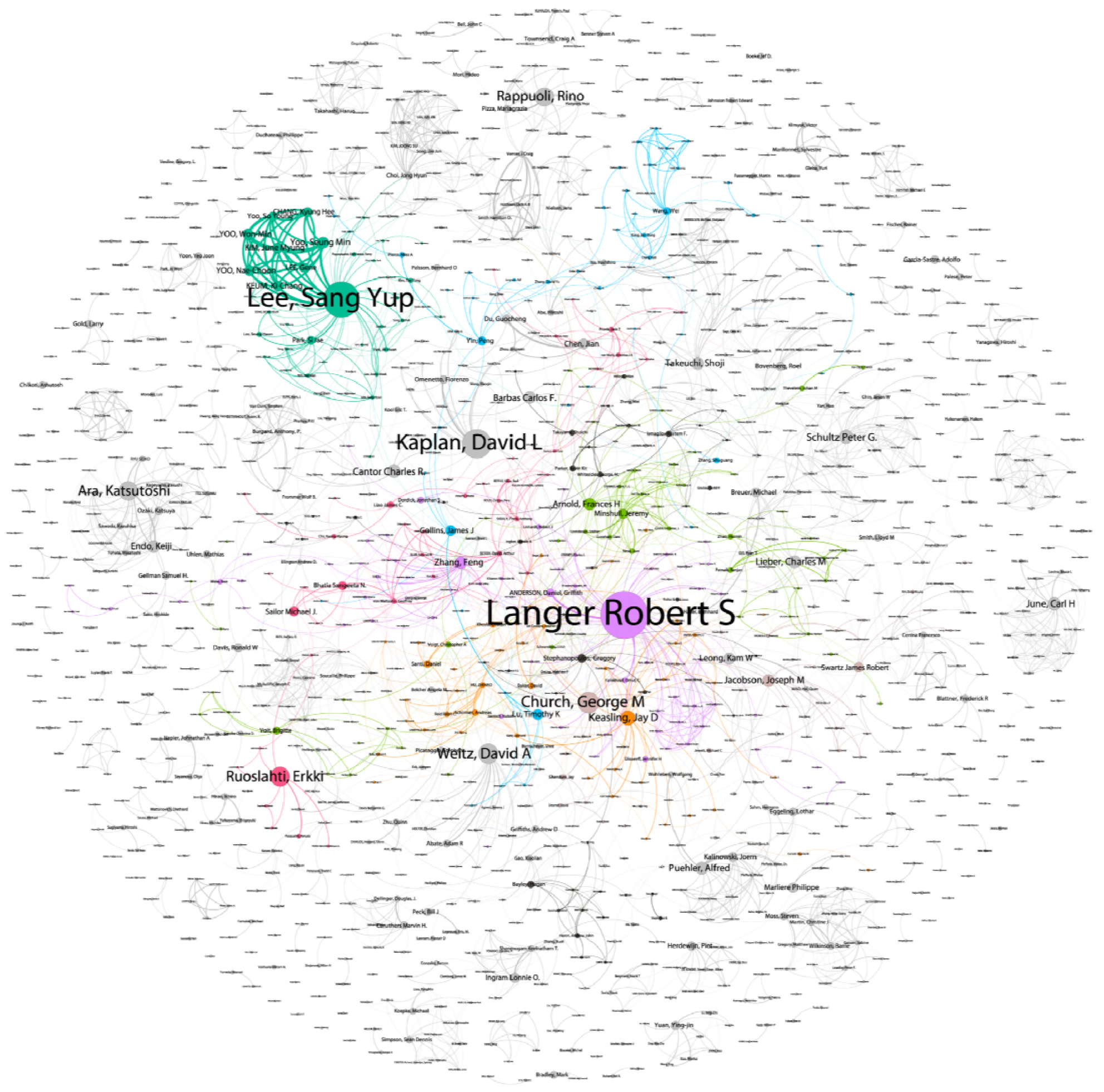
Inventor Communities.

As discussed above in relation to the use of counts of coauthors as inventors, Robert Langer at MIT has a very low score (1) in terms of the number of publications containing the baseline terms, but ranks top in terms of the number of patent filings, family members and scores highly for the number of intersecting authors (19) who are also inventors. Robert Langer also has the highest citing patent score in terms of impact within the patent technology space. Robert Langer is therefore a logical place to start in exploring individual networks. Figure 23 sets out the network of inventors for Robert Langer based on their appearance in five or more patent filings. The network is coloured so that authors appearing in the literature dataset appear in green and those outside it in pink. The network has been truncated to facilitate display.

**Figure 23:**
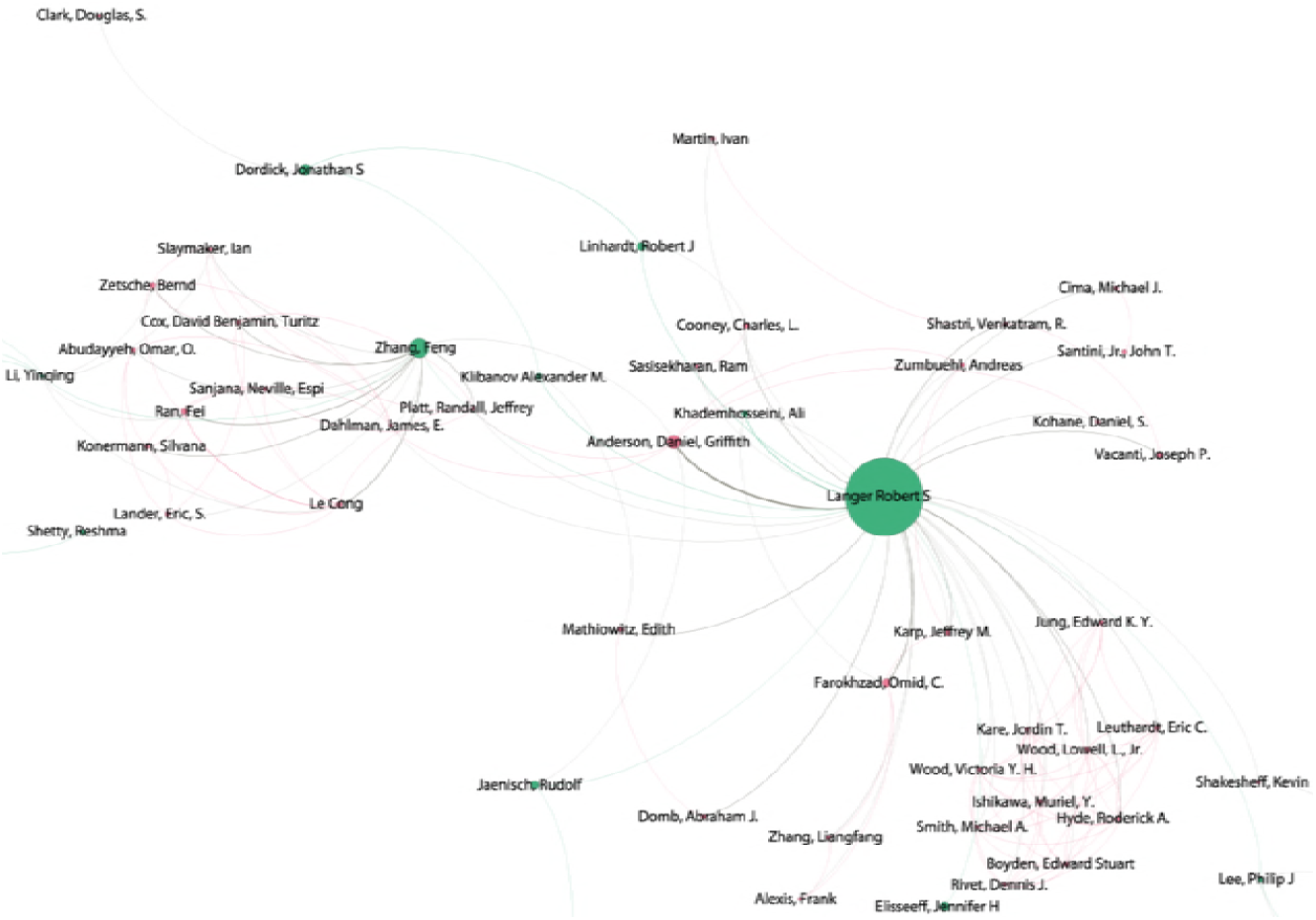
The Langer Network.

The Langer network is particularly significant with respect to synthetic biology for the link to Feng Zhang at MIT in connection with CRISPR genome editing. Robert Langer is a co-inventor with Feng Zhang for the “Delivery Use and Therapeutic Applications of the CRISPR CAS Systems and Compositions for Targeting Disorders and Diseases using Particle Delivery Components” (IN201617023305A, US20150232883A1, SG11201604406A1). Robert Langer is also a co-inventor with coauthor Ali Khademhosseini on applications relating to amplification of cell populations from embryonic stem cells (US20070042491A1), cell laden hydrogels (US20080193536A1) and microfluidic arrays (US20070266801A1). With recent Nobel prize winner Frances Arnold at Caltech (not shown), Robert Langer was also a co-inventor on an application for Metalloprotein MRI contrast agents (WO2009032160A2). Community networks also include indirect relationships stemming from the focus researcher. In the case of the Langer network, Yinqing Li is a coauthor inventor with Feng Zhang who has also submitted patent applications with Ron Weiss for Layered Transcriptional Circuitry using CRISPR Systems (US20160326546A1) and novel recombinases and target sequences (US20170211061A1). Feng Zhang also links with Cong Le, whose work has focused on genome engineering with CRISPR/CAS systems, but falls outside our baseline literature dataset [61]. Feng Zhang and Le Cong are co-inventors on 10 filings relating to CRISPR within the synthetic biology landscape (e.g. in connection with CAS9 orthologs in WO2016205759A1).

The second major network on the inventor level, measured on the number of first filings, is Sang Yup Lee at the Korea Advanced Institute of Science and Technology (KAIST). Figure 24 displays the main elements of this community.

**Figure 24:**
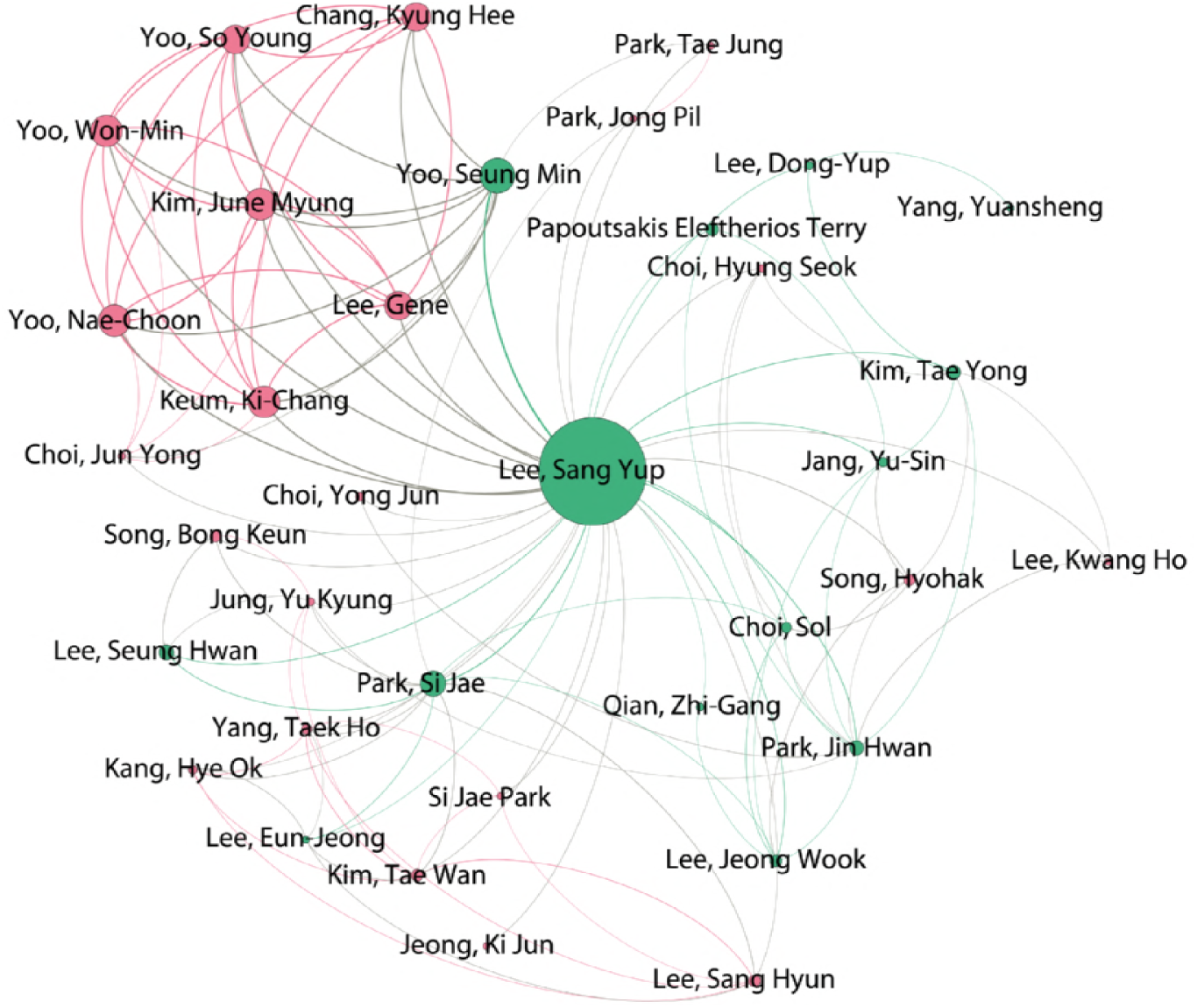
The Sang Yup Lee Network.

Sang Yup Lee’s patent portfolio consists of a cluster of filings in the mid-2000s for DNA Chips for detecting a range of pathogenic bacteria, such as *Escherichia coli, Staphylococcus aureus*, and *Clostridium difficile* among others (e.g. KR2007069676A, WO2008084889A1, KR2005084549A). Highly cited patent families include Novel rumen bacteria variants for preparing succinic acid (WO2005052135A1) and gene encoding formate dehydrogenases for preparing succinic acid (US7241594B2) along with mutants for producing large quantities of butanediol using microorganisms producing succinate (KR2007096348A)[62]. In other filings, reflecting wider interest across the landscape and linking to clothing, Sang Yup Lee has also filed in relation to recombinant spider silk (WO2011112046A2) [63]. More recent work has focused on CRISPR/CAS9 engineering of Actinomycetal genomes, notably those where gene editing is difficult such as streptomycetes (WO2016150855A1). Streptomycetes are soil bacteria that have historically been the world’s single major source of antibiotics [64].

Figure 25 displays the main network for David Kaplan at Tufts College. This network reveals highly cited patent filings for silk biomaterials (US20050260706A1), antioxidant functionalised polymers (WO2004050795A2), biocompatible scaffolds and adipose derived stem cells (WO2007103442A1), and silk fibrion microspheres using lipid vesicles for controlled release of therapeutic agents (WO2008118133A2). Filings with co-author inventor Charles Cantor in the early and late 1990s include biotin-binding containment systems and (WO1996034954A2) along with compositions and methods for controlling genetically engineered organisms through cell suicide involving the expression of the lethal Streptomyces avidinii streptavidin gene (WO1999040179A1). More recent filings include 3D printing of biopolymer based inks (WO2016019078A1), biodegradeable electronic devices (WO2008085904A1), silk based adhesives (WO2017095782A1), electroactive scaffolds (WO2017011452A1) and shape memory silk materials (WO2016145281A1).

**Figure 25:**
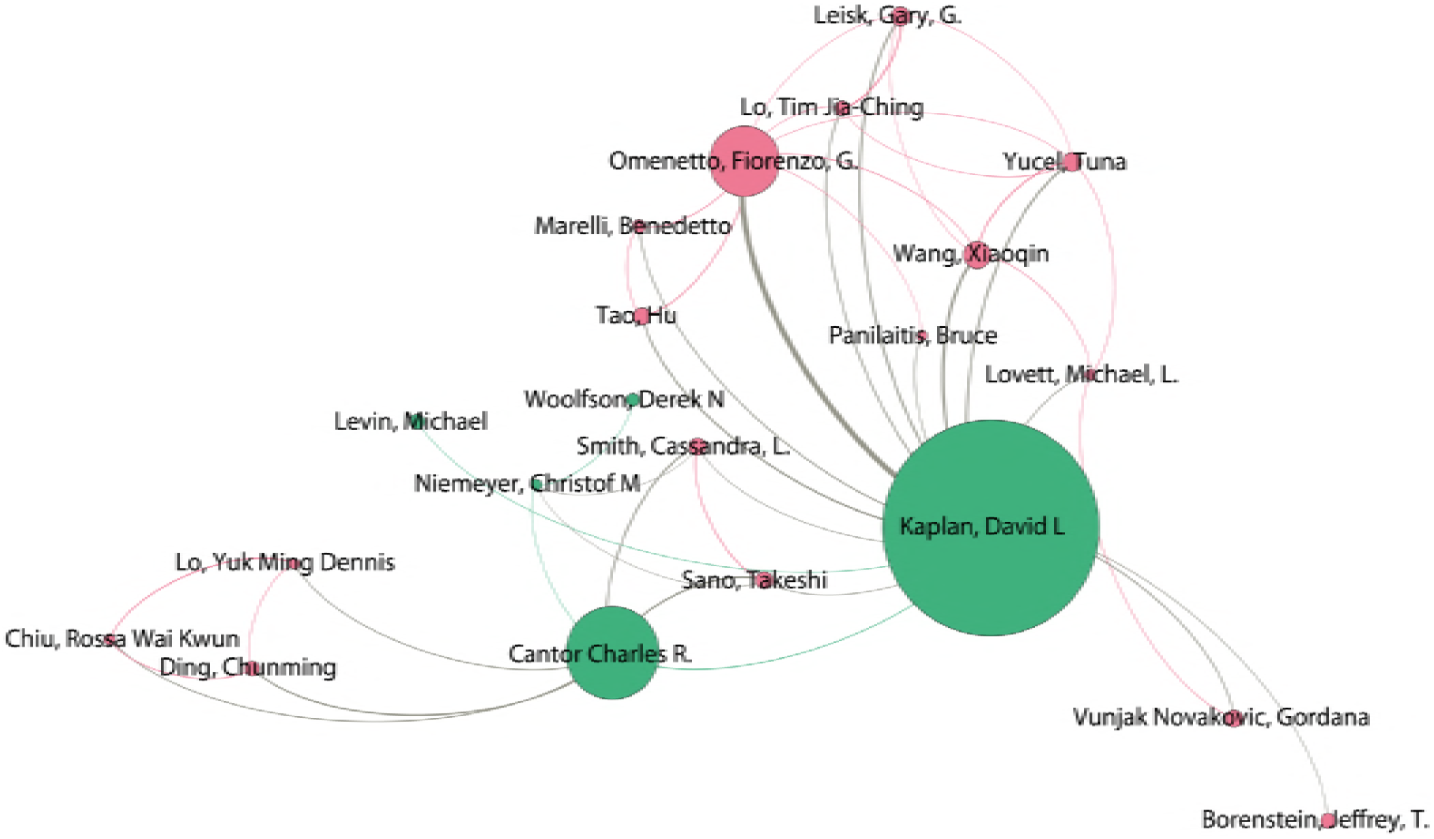
The Kaplan Network.

Figure 26 displays the network focusing on George Church at Harvard and Joseph Jacobson at MIT. Patent activity by George Church with significant impact dates back to the mid-1980s and includes work on multiplex analysis of DNA (EP303459A2), replica amplification of nucleic acid arrays (US6432360B1) and surface bound double stranded nucleic acid and protein arrays (WO1999007888A1, WO1999019510A1) in the late 1990s. In the mid-2000s activity focused on wobble sequencing (WO2006073504A2), methods for assembly of high fidelity synthetic polynucleotides (WO2006044956A1), nanogrid rolling circle dna sequencing (WO2007120208A2), and hierarchical assembly methods for genome engineering (US2007004041A1). More recent activity includes RNA guided systems for probing and mapping of nucleic acids (CA2958292A1), and systems and methods for processing spatially related sequence data from a sequencing device (WO2017079382A1). In 2016 a set of 9 filings focused on issues such as enzymatic nucleic acid synthesis (WO2017176541A1), cell free enzyme discovery and optimization (WO2017155945A1), molecular recording by CRISPR-CAS system (WO2017142999A2), nuclease mediated gene editing in stem cells (WO2017184674A1), rule based genome design (WO2017218727A1) and recombinase genome editing (WO2017184227A2).

**Figure 26:**
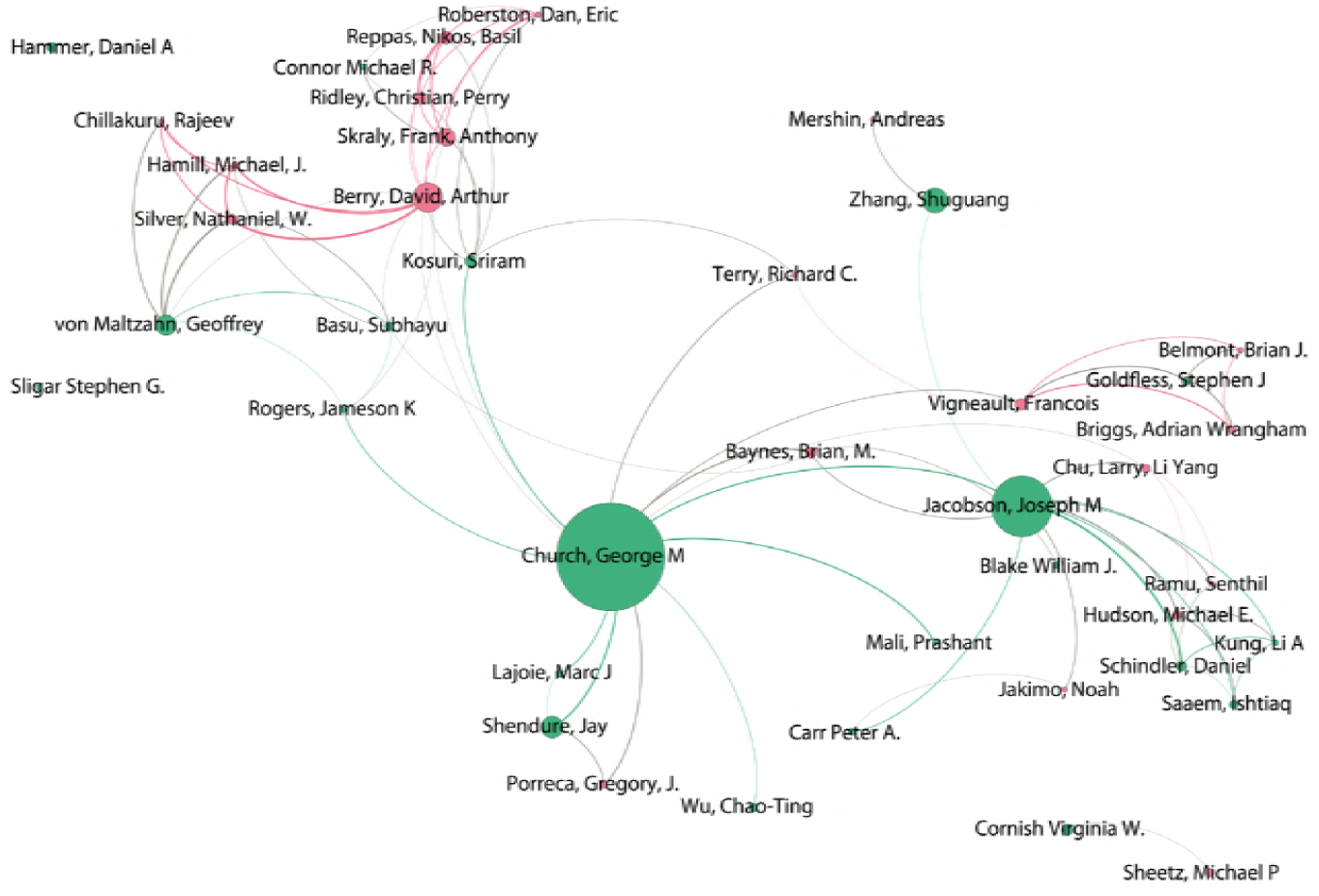
The Church/Jacobson Network.

Looking beyond the emphasis on whole genome engineering and genome editing in this portfolio, George Church has also filed five joint patent applications in our dataset with Joseph Jacobson at MIT on subjects such as engineered metabolic pathways (WO2008127283A2), aptamer production (WO2007136833A2), and gene regulation (US20120315670A1). As Figure 26 reveals Joseph Jacobson could be treated as a separate network and has contributed to scientific research and innovation in at least two areas. The first of these is the development of microencapsulated electrophoretic displays (E Ink) that are now widely used in electronic reader devices. In practice, E Ink devices arising from this work form a distinct landscape (see for example, US6262706B1, US6512354B2, US6480182B2) involving distinct inventors and is not considered in connection with synthetic biology. Patent applications with other authors of articles on synthetic biology include filings on high fidelity assembly of nucleic acids (WO2013032850A2), multiplex nucleic acid synthesis (US20150376602A1), *in situ* nucleic acid synthesis (WO2011150168A1), preparative *in vitro* cloning (WO2012174337A1), cell based genomic recorded accumulative memory (known as geRAM or Genomically Encoded Memory GEM) (US20150133315A1) and population hastened assembly genetic engineering for continuous genome recording (US20160244784A1). Outside the coauthor network the increasing influence of CRISPR/Cas9 in the synthetic biology landscape is perhaps reflected in a filing for programmable Cas9 biomolecular circuitry with RNA input (WO2015168404A1).

Figure 27 displays the network focusing on Rino Rappuoli, formerly at Chiron and Novartis. Research by Rappuoli [65] and close associate Mariagrazia Pizza [66] has focused on the use of rational design of molecules and reverse vaccinology. This has led to globally significant developments in connection with diphtheria and in carrier development (specifically the use of CRM197 as a carrier in *H. influenzae*, pneumococcus and meningococcus vaccines), and vaccines for pertussis (whooping cough) using site directed mutagenesis, and meningococcus B (meningitis) using reverse vaccinology. The rise of reverse vaccinology was facilitated by a collaboration with Craig Venter and colleagues to sequence the genome of Neisseria meningitides subtype B [65].

**Figure 27:**
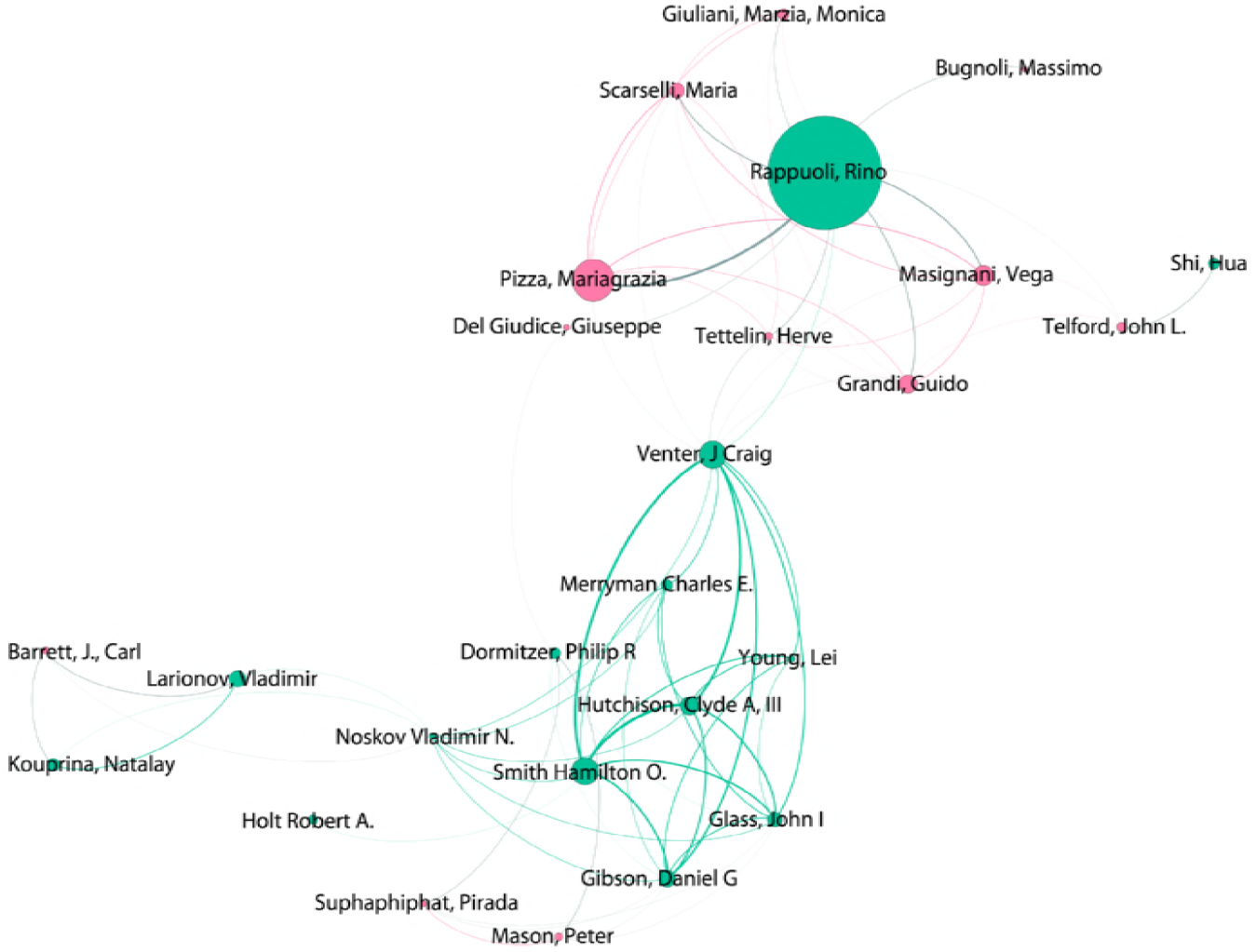
The Rappuoli/Venter Group Network.

As Figure 27 reveals work by Rappuoli and close associate Mariagrazia Pizza links to a cluster of researchers who form part what is now the J. Craig Venter Institute and Synthetic Genomics with Nobel winner Hamilton Smith, Clyde Hutchinson, John Glass and Daniel Gibson as prominent coauthor inventors. In the mid-1990s filings in connection with whole genomes (such as *Haemophilus influenzae, Mycoplasma genitalium, Methanococcus jannaschii* and *Neisseria meningitidis*) attracted significant attention [67]. However, whole genome sequence patent applications in the United States and Europe have been regarded as lacking unity of invention, because a whole genome sequence is not regarded as a single invention, and have been restricted to fragments upon examination [67]. In light of the collaboration with the Rappuoli group, whole genome claims arising from research by the Venter group might reasonably be seen as defensive in nature and perhaps as serving to showcase their research to an international public.

More recent activity in the mid-2000s is explicitly associated with the emergence of synthetic biology or synthetic genomics. The initial cluster with earliest filing dates in 2005 focused on a minimal bacterial genome (AU2006303957A1), synthetic genomes (US20070264688A1), methods for *in vitro* recombination (US20070037196A1), and genome installation (transplantation of a genome into an empty cell) (US20070269862A1). A second cluster of filings in 2007 focused on the assembly of large nucleic acids, methods for cloning and manipulating genomes, further elaboration on genome installation and a system and method for producing synthetic microorganisms translating nonstandard amino acids (AU2008310912A1). These were followed in early 2008 by a filing on methods for in vitro joining and combinatorial assembly of nucleic acids (US20120053087A1). A 2009 filing linked to the earlier filings addressed methods for cloning and manipulating genomes (US2016177322A1). In 2012 a small cluster of filings addressed: synthesis of error minimized nucleic acids (CA2862149A1), crowding agent induced DNA transfer into a host cell (US20140179001A1), polyethylene glycol (PEG) mediated assembly of nucleic acids (US20140308710A1). An additional filing in 2012 involves a digital to biological converter for transmitting sequence information from a remote location to initiate synthesis of a biological entity (US9718060B2). The aim of this application is to transmit the data necessary to produce a vaccine to a remote location as part of rapid response to biological threats. In 2013 this was followed by an application for influenza virus reassortment focusing on creating a virosomal vaccine (WO2014115104A1) and a separate application for the transplantation of nucleic acids into cells using microfluidic chambers (US9834747B2). A 2017 filing focuses on the generation of synthetic genomes (WO2017165565A1).

Figure 28 displays the Jay Keasling network. Jay Keasling at the University of California Berkeley and Amyris Inc is one of the most prominent actors in both the scientific and patent landscapes for synthetic biology with 50 publications under the baseline definition and 49 patent families. Here we focus on a brief overview of the portfolio using highly cited families. Patent activity within our dataset can be traced to a 2001 filing for biosynthesis of isopentenyl pyrophosphate and derivative isoprenoids (US20030148479A1) with additional filings on isoprenoids in 2004 including enhancing production (US20060079476A1) and in 2005 on polynucleotides encoding isoprenoid modifying enzymes (CA2613469A1). Other highly cited filings focus on diacids through polyketide synthesis (WO2009121066A1) and host cells for producing diacids (WO2012071439A1). The highest cited filing in the Keasling portfolio, with George Church from Harvard as a co-inventor, is for production of fatty acids and derivatives that addresses the problem of cracking biocrude, hydrocarbon feedstock produced by a microorganism and producing a biofuel (EP2395074A1).

**Figure 28:**
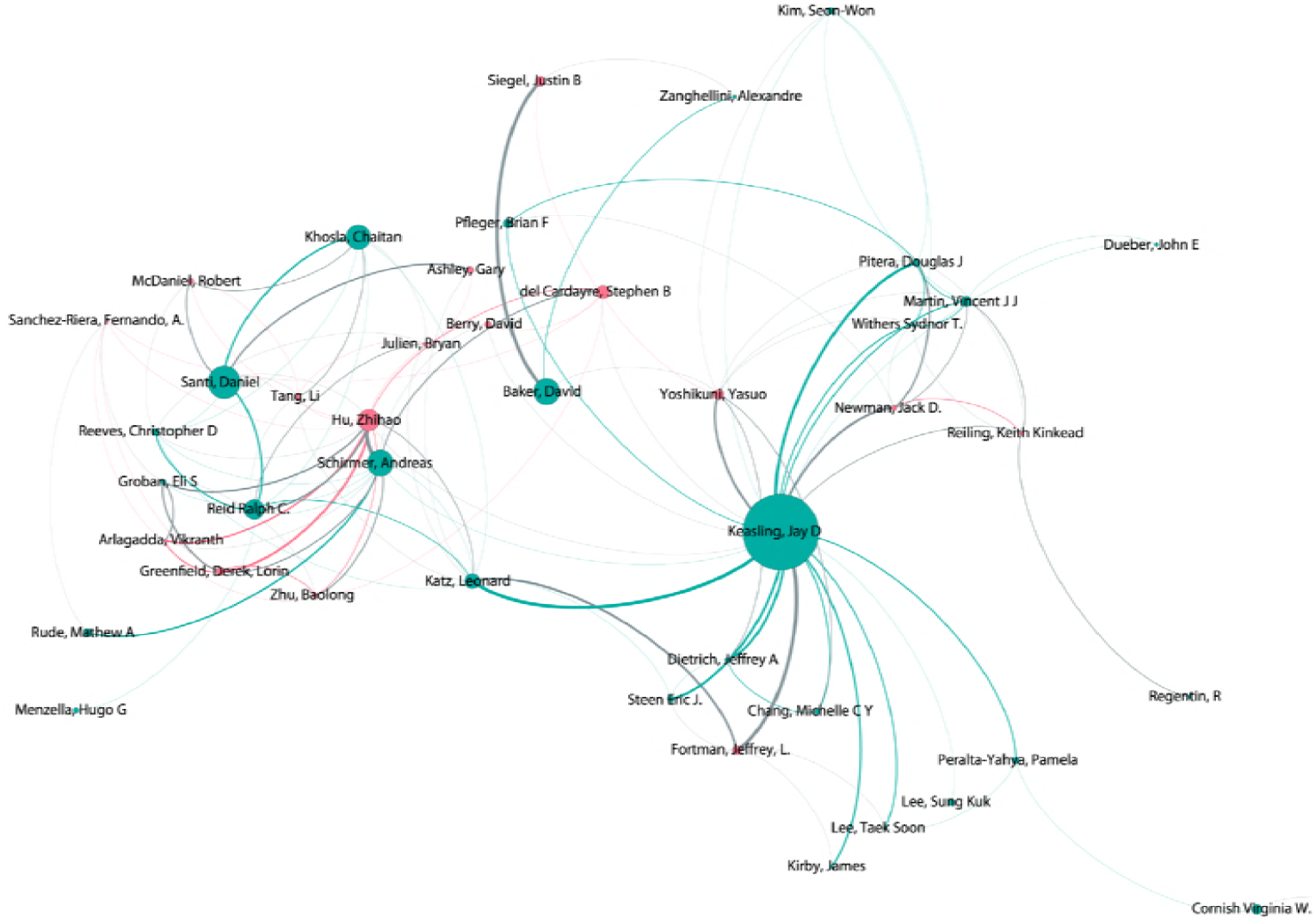
The Keasling Network.

Jay Keasling’s work is closely associated with synthetic artemisinin for the treatment of malaria [68,69]. Interest in this field is reflected in the 2001 filing on biosynthesis of isopentenyl pyrophosphate highlighted above and the synthesis of artemisinic epoxide (WO2008073498A2). Amyris Inc, founded by Jay Keasling, has also filed for conversion of amorpha-4, 11-diene to artemisinin and artemisinin precursors (CA2609331A1). The licensing of the production of semi-synthetic artemisinin to Sanofi led to considerable interest in the potential for synthetic biology to address supply side issues for natural artemisinin while raising concerns that synthetic production could impact on the livelihoods of producers of the natural version [70–72]. However, in 2016 it was reported that Sanofi would not proceed with production and was selling its production facility [73]. At the time of writing recent reports suggest that production is under way and that more efficient routes to semi-synthetic synthesis have been identified [74].

Our focus so far has been on the exploration of top inventors using a combination of counts by patent families and co-author counts. The use of coauthor counts links to the use of a range of centrality measures in social network analysis [75]. For our purposes the most important is betweeness centrality that ranks nodes in a network based on the measurement of the shortest paths between nodes [76]. That is, betweeness centrality identifies nodes that are the most important bridges between others in a network. Figure 29 reproduces Figure 22 but resizes the nodes by betweeness centrality. Minor adjustments have been made to node position for label visibility and edges may intersect with unrelated nodes.

**Figure 29:**
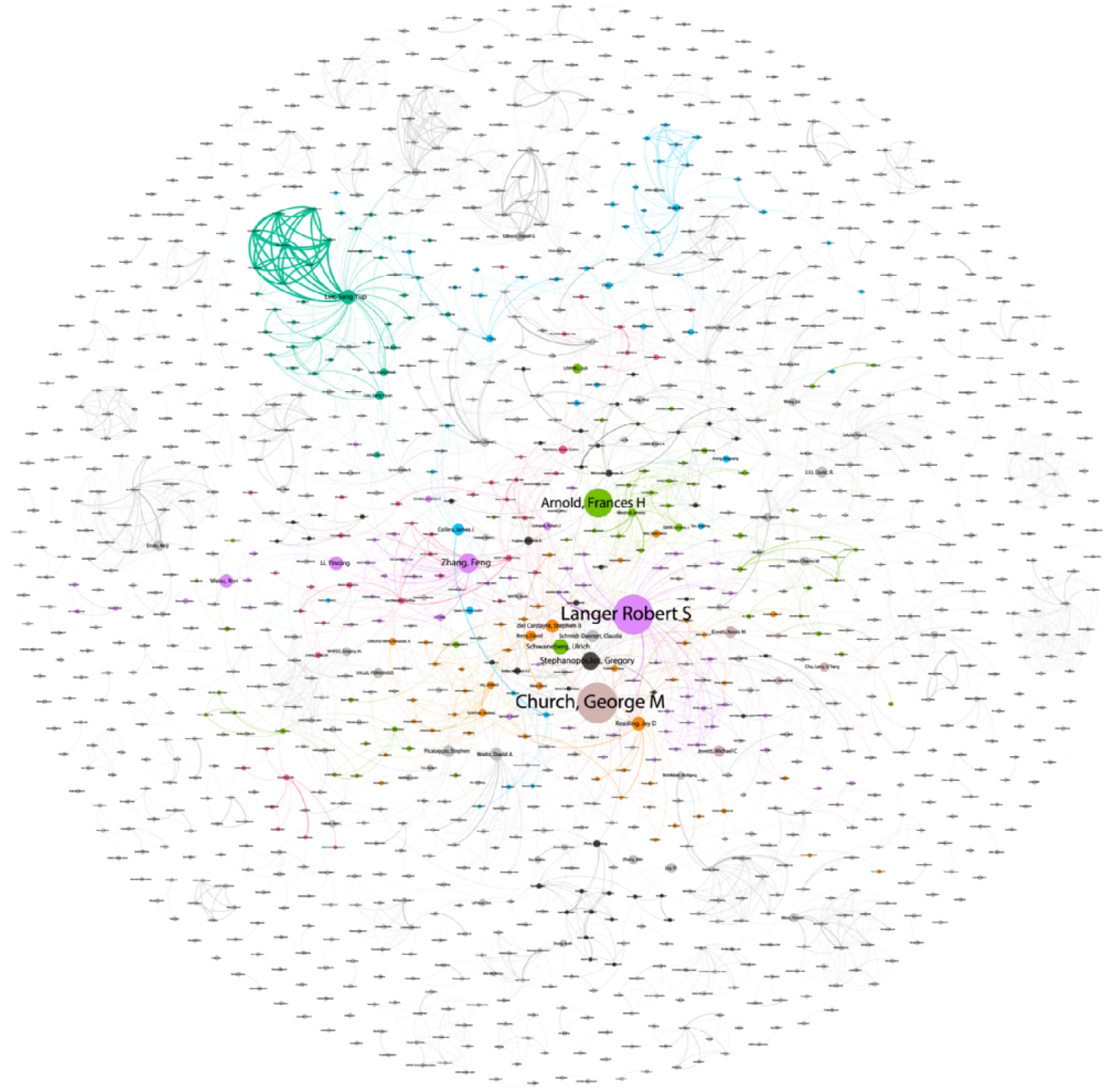
Central Actors in the Inventor Network by Betweeness Centrality.

When viewed in terms of betweeness centrality scores, we observe the prominent position of the joint winner of the 2018 Nobel prize for chemistry Francis Arnold at Caltech for work on the directed evolution of enzymes [77].

In the case of Francis Arnold, we observe highly cited filings for a microfabricated cell sorter in the mid 1990s (CA2333201A1) along with a method for evolving polynucleotides with desired properties (US6537746B2). A cluster of filings between 1999 and 2001 focused on directed evolution including for the directed evolution of biosynthetic and biodegradation pathways (WO2001042455A1), computationally targeted evolutionary design (WO2001061344A1), and directed evolution of oxidase enzymes (WO2001088110A1) along with a screening methods for the discovery and directed evolution of oxygenase enzymes (WO2001077368A1) followed by work on Cytochrome P450 oxygenases (WO2003008563A2, WO2002083868A2). A more recent cluster of filings between 2009 and 2010 focused on polypeptides with cellulase activity (WO2010118058A2), regioselective alkane hydroxylation (US8309333B1, US8361769B1) and engineered microorganisms with metabolic pathways capable of producing C3-C5 alcohols under aerobic or anaerobic conditions (CA2779262A1). The most recent filings in our dataset from 2013 and 2014 provide an endoglucanase with enhanced thermostability (US20140308713A1) and the sulfimidation or sulfoximidation of organic molecules (US9399762B2).

The networks discussed above form part of a wider inventor landscape of approximately 223 communities or clusters involving 2,450 researchers publishing on synthetic biology who are engaged in patent activity, their associates and others active in the field of synthetic biology. As we have seen the main clusters revolve around the MIT - Harvard - University of California axis. These networks form complex webs of collaboration characterised by flows of funding, training, mentorship, patronage, friendships, competition and knowledge production directed towards commercialisation as an offshoot or objective of research. These networks also form the basis for the organisational relationships considered above in terms of partnerships with major commercial companies and the establishment of spin off companies.

As we have seen patent activity at the level of individual researchers can be approached using a variety of measures. Each of those measures provides a different perspective on the wider landscape. In particular, we have highlighted that care is required in the assessment of the presence of senior researchers with large patent portfolios but limited publications in an emerging field. In the case of the J Craig Venter Institute, in our view, the orientation of their research cannot adequately be understood if the connection with the work of Rino Rappuoli is simply excluded. Similarly, as we have seen in the case of Francis Arnold, the influence of researchers within the patent landscape is not reflected at the level of simple counts. As such one important insight from this approach is that a range of measures are desirable when conducting analysis at the researcher level.

### The Impacts of Patent Activity

Patents are a common focus for public debate and controversy because they are visible indicators of commercial developments in science and technology. In closing this analysis of the patent landscape for synthetic biology we will briefly consider the impacts of patent activity in synthetic biology. This is likely to involve four main issues.

First, patents have a role to play within biotechnology in raising finance and facilitating partnerships with companies. Within academia patents are also increasingly seen as an indicator or achievement and may in some cases generate significant revenue for University employers. However, patent claims by researchers in synthetic biology may potentially have impacts on the openness of synthetic biology as an emerging field [8,19]. The extent to which this is presently a problem within the community is an open question. The data provided with this article provides a starting point for more in depth evidence based discussion on patent activity within the community.

A second issue is the wider impact of patent activity on the production of new and useful products, methods, or processes that enter the market. Within the economics literature patent protection is commonly linked with Foreign Direct Investment (FDI) and creating an environment conducive to investment and corresponding knowledge spillovers [78]. Conversely, patent protection may also be associated with license based rent extraction, as in the historic patent controversies on Taq DNA polymerase and BRCA1 and 2, and trolling behaviour. Addressing these issues is not straightforward, particularly in circumstances where one patent is commonly not equivalent to a single ‘product’ as such but forms part of a product made up of many elements.

A third issue, of direct relevance to debates under the United Nations Convention on Biological Diversity, concerns the actual or potential impacts of synthetic biology upon the environment and the welfare of indigenous peoples and local communities. Once again, we will not address these issues directly. However, the evidence provided with this article can support evidence based deliberation on issues such as genome editing, gene drives and the role of digital sequence information in synthetic biology.

The citing patent landscape for synthetic biology also has an important role to play in considering these issues because it provides additional data on actors within the patent system who are being affected by synthetic biology and provides insights into emerging developments such as genome editing and gene drives.

The citing landscape for synthetic biology consists of later patent filings with claims that are limited by the scope of claims in the core landscape. The citing landscape is expected to include filings by others working in the field who fall outside the baseline search query and companies and organisations whose property claims are affected by filings from core group. The citing landscape was prepared by removing all filings that already appear in the core landscape. This reduced the citing landscape to 33,889 families and 499,284 family members. After name cleaning we identified approximately 13,257 applicants, including individuals, in the citing landscape in the period to December 2017.

In approaching the citing landscape it is important to emphasise that patent citations may originate from across the spectrum of technology within the patent system. That is, a minor technical feature within a document may lead to a citation in a later filing. As such, the citing landscape will include data that is both directly relevant to synthetic biology and data that is irrelevant to the emerging field.

Figure 30 displays the top ranking applicants in the core dataset based on counts of first filings. Colours denote organisations that already appear in the core landscape and those falling outside the core landscape. A total of 983 applicants already appeared in the core landscape with the remaining 12,274 being new. Of the new applicants only 2,432 organisations had more than one filing affected by a filing from the core landscape.

**Figure 30:**
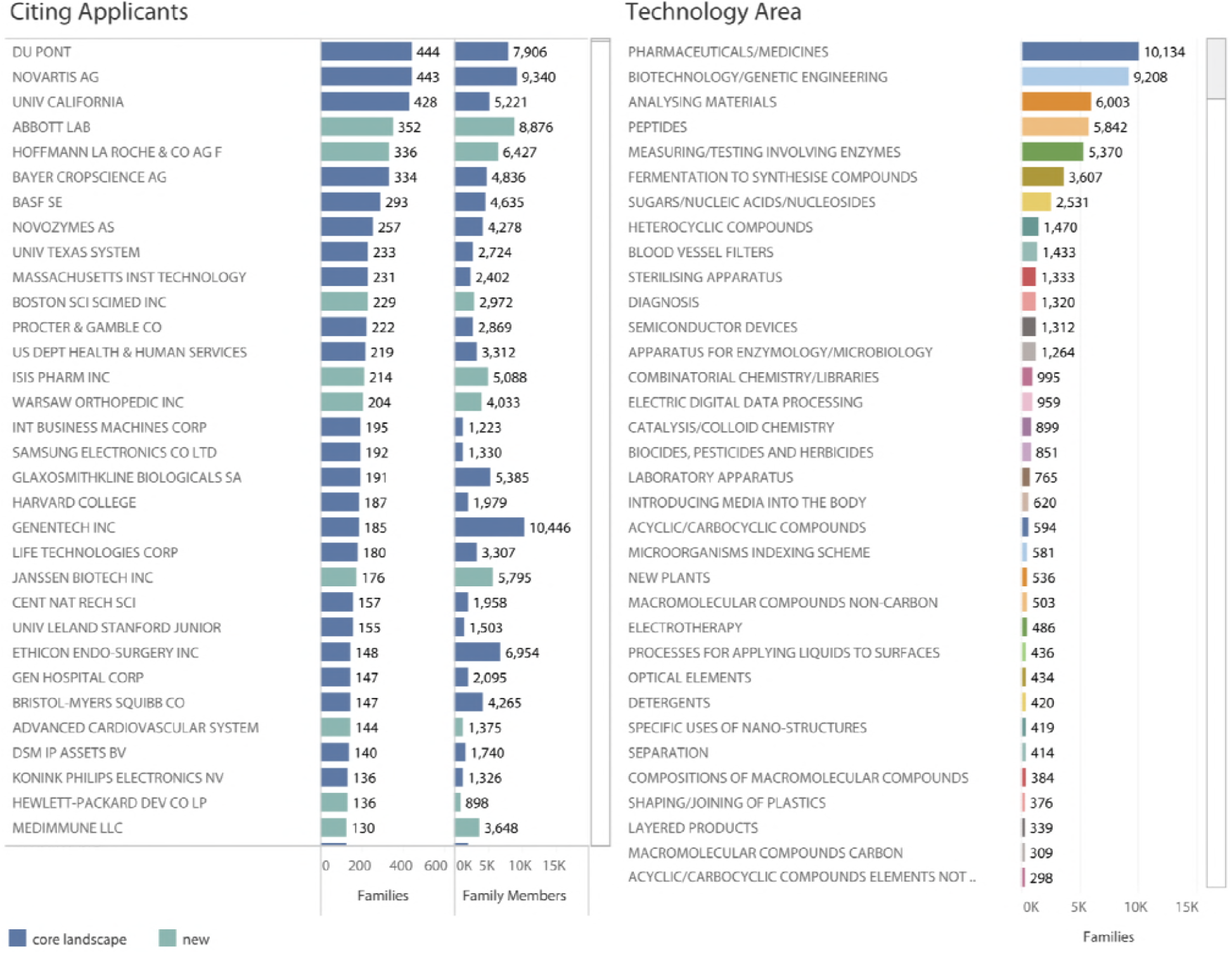
Citing Landscape Applicants and Technology Areas.

Figure 30 reveals that those companies and research organisations that are being affected by patent activity in synthetic biology are commonly already located inside the core landscape dataset. These will be cases where applicants are being affected by competing claims from other applicants within the core landscape.

In considering Figure 30 note that while top ranking universities from the core landscape appear in the top applicants in the citing landscape, companies are becoming increasingly prominent. In addition we also observe companies situated outside the patent landscape beginning to enter the picture such as Abbot Labs and Hoffmann La Roche. Turning to technology areas based on IPC sub-classes, we observe the dominance of pharmaceuticals (compared with the core landscape) followed by biotechnology related activity. As this makes clear both the core landscape for synthetic biology and the citing landscape are firmly situated in the field of biotechnology and genetic engineering.

Taking into account the simple baseline query used to generate the core landscape we believe that the wider landscape of researchers working in synthetic biology in areas such as CRISPR Cas9 and synthetic biology companies are likely to be situated in the citing landscape. Full exploration of this possibility is beyond the scope of this article. However, we can illustrate this for CRISPR Cas9.

As noted above CRISPR Cas9 is an increasing feature of the patent landscape for synthetic biology and we identified approximately 211 families and 1835 family members worldwide referencing CRISPR or Cas in the title, abstracts or claims. As discussed above in connection with the Langer network, for CRISPR data the lead inventor in the core landscape is Feng Zhang at MIT with 53 families followed by George Church (6) and Kevin Esvelt (5) whose work on CRISPR Cas9 is associated with gene drives [79,80]. CRISPR Cas9 also features in the core landscape in work at UC Berkeley by James Cate Doudna with Owen Ryan inside our author inventor set (WO2015138855A1) and Jennifer Doudna at UC Berkeley (WO2017106569A1 on modified site-directed modifying polypeptides) who falls within the keyword dataset on the term synthetic genome. The presence of these actors in the core dataset is relevant because of the litigation between the Broad Institute and UC Berkeley over CRIPSR Cas9 patent filings [81]. The two key patent documents in this dispute as reported by Egelie et. al 2016 are US8697359 and US20140068797 (CA2872241A1) from UC Berkeley [81]. Both documents appear in the core landscape with the UC Berkeley filing appearing in the core set by virtue of Wendell Lim at UCSF being listed as a co-inventor. The third party in the dispute is Vilnius University and researcher Virginijus Siksnys with a family in the citing landscape for RNA-Directed cleavage by the Cas9-CrRNA complex (EP2828386A1) and rapid characterization of CAS Endonuclease Systems (WO2016186946A1).

The at times heated dispute over CRISPR Cas9 has recently been concluded in favour of the Broad Institute, which used expedited examination before the USPTO to move ahead of potential competitors to good effect, pending a possible appeal to the US Supreme Court [82–84].

This dispute is however only part of the story. Within the citing landscape we identified an initial 151 families referencing CRISPR or Cas in the title, abstracts or claims, with leading applicants including Du Pont, Sangamo Biosciences, Regeneron Pharma, Novartis, Caribou Biosciences, Dow Agrosciences, CRISPR therapeutics and Editas Medicine. While meriting more detailed analysis, the citing landscape includes the main actors identified in the CRISPR landscape by Egelie et. al. 2016 [81]. Furthermore, the citing landscape also captures emerging activity directed towards CRISPR 2.0 such as work by David Liu at Harvard in connection with target sites for Cas9 nucleases (US9163284B2), Cas9-Foki fusion proteins (US9322037B2) and evolution of site specific recombinases (WO2017015545A1).

Further exploration of the citing landscape is beyond the scope of this article. However, as the discussion above suggests the method successfully captures key developments within the synthetic biology community network itself while the citation system will capture additional activity. As noted above, citation data will also include results that are irrelevant to synthetic biology. However, a modular approach to navigating citation networks using key words to target specific areas of interest as described by Arora et. al. 2012 is likely to open up more detailed landscapes and facilitate tracking of emerging developments [85,86].

## Conclusion

This article has aimed to contribute to mapping the international patent landscape for synthetic biology as an emerging area of science and technology. Starting with a baseline key word query specific to synthetic biology we mapped authors of articles on synthetic biology into the patent system. We then used the patent citation system to capture additional actors involved in synthetic biology and a keyword dataset to capture additional records.

The concentric search model is intended to help overcome the challenges presented by keyword based approaches for mapping emerging areas of science and technology in cases where a new field is emerging *within* existing fields using very similar terminology. A particular strength of the approach we adopted is that by focusing on people it favours precision at the level of the identification of the work of individual researchers. The method then seeks to complete the landscape by expanding data capture using citations and keywords. As we have seen by focusing on what researchers are actually doing in the patent system we are able to engage in detailed exploration of networks of research and innovation. We are also able to accurately address emerging developments in a field, such as the use of CRISPR Cas9, genome editing and gene drives, by searching within community networks and citation networks.

The method we described involves the use of both scientific publication data and full text patent data. While commercial services such as Clarivate Analytics offer high quality data and important additional value, the ability to conduct patent research at scale is constrained by issues of data access. That situation is beginning to change in the case of the scientific literature as a result of initiatives such as Microsoft Academic Graph and Open Academic Graph. In the case of patent data the PatentsView service established by the USPTO now makes the full US collection available for research in a convenient form. Other services such as the open access Lens Patent database and WIPO Patentscope offer additional value. Increasing access to patent data promises to overcome some of the limitations confronting earlier research and at the same time to promote greater methodological transparency [29]. The World Intellectual Property Organization is actively seeking to promote greater methodological rigour in patent analytics through a series of wide ranging patent landscape reports at varying scales [87]. However, a key constraint in making use of larger scale resources is the ability to address the name disambiguation problem both within and across data sources. As discussed by Morrison et. al. 2017 a range of programmatic approaches focusing on machine learning are now available [88]. However, many of these approaches address specific problems rather than presenting generalisable solutions [88]. In recognition of the real world complexity of name matching and disambiguation across data sources, we provide a tagged dataset to contribute to further methodological development in machine learning and related approaches.

The people centred approach to landscape analysis presented in this article is also relevant to debates on intellectual property within the synthetic biology community. Our research reveals that approximately 23% of researchers publishing scientific research in the field of synthetic biology are also active in the patent system. That is the majority of researchers publishing using terms such as synthetic biology or synthetic genomics are not engaged in patent activity. As we have seen those researchers who are engaged in patent activity form distinct networks, predominantly, but not exclusively, concentrated in the United States. The data we provide is intended to assist with evidence based discussion of the implications of patent activity within the synthetic biology community in areas such as standards, or the uses of tools such as CRISPR.

The outcomes of the present research are also relevant to debates on synthetic biology, and emerging debates on gene drives and digital sequence information (sequence data), at the Convention on Biological Diversity and under its Nagoya Protocol in three ways [89]. First, it provides actual evidence of the main actors involved in patent activity for synthetic biology. Second, delegations have struggled to understand the distinction between genetic engineering and Living Modified Organisms as addressed under the Cartagena Protocol on Biosafety and synthetic biology. Approaching synthetic biology as the synthetic phase of genetic engineering that is reflected in continuities and departures from genetic engineering may assist with enhancing understanding. Specifically, it may assist with developing a longer term policy perspective on this field and place recent developments such as genome editing and gene drives in their wider context. By focusing on evidence, and identifying lead actors and networks, Parties to the Convention will also be in a better position to assess the interests at stake and to consider the positive and negative implications of developments in this field for the objectives of the Convention with respect to conservation, sustainable use and fair and equitable benefit-sharing arising from the utilisation of genetic resources.

The focus on people within this article also directs our attention to issues relating to social and environmental responsibility and future developments in this field. The growing institutionalisation of synthetic biology and establishment of undergraduate and doctoral programmes suggests a shift to increasing professionalisation within the field. Within the context of the Convention this allows us to ask questions about the standards set by funding agencies and the training those entering this field receive with respect to social and environmental issues. In the UK, while modest in scale, investments in Responsible Research and Innovation form part of research investments in synthetic biology that bring social scientists and synthetic biology researchers together with varying degrees of mutual comprehension [90–92]. However, this raises the wider question of whether additional guidance to funding agencies and the synthetic biology community on social and environmental issues relevant to the implementation of the Convention might form part of an appropriate way forward.

It remains to be seen what Parties to the Convention will decide in addressing synthetic biology. However, recent years have witnessed increasing efforts within the social science community and the synthetic biology community itself to improve the evidence base to inform social, economic and environmental debates on synthetic biology. The present work aims to contribute to strengthening evidence based debate and to provide a platform for further methodological innovation.

## 1 Acknowledgements

Paul Oldham thanks Philip Shapira for critical comments on an earlier version of the manuscript and the members of the Responsible Research and Innovation Group at SYNBIOCHEM Manchester University for their support and encouragement while completing the research.

